# Growth in Low Carbon Conditions Reveals Amino-Acid-Coupled Iron Uptake

**DOI:** 10.1101/2025.01.15.633289

**Authors:** Juanita Lara-Gutiérrez, Jen Nguyen, Matthew R. McIlvin, Ichiko Sugiyama, Zachary Landry, Uria Alcolombri, Johannes M. Keegstra, Sammy Pontrelli, Joaquín Jiménez-Martínez, Uwe Sauer, Terence Hwa, Mak A. Saito, Roman Stocker

## Abstract

Bacteria in nature encounter substrates at widely varying concentrations, yet studies of bacterial physiology have focused more on nutrient type than concentration, partly due to challenges in maintaining low concentrations. We developed a Millifluidic Continuous Culture Device (MCCD) to culture bacteria under precisely controlled nutrient conditions, including very low concentrations, in a manner suitable for proteomic analysis. Using the MCCD, we cultured *Escherichia coli* with a mixture of amino acids as the sole carbon source at three concentrations supporting growth rates spanning a fivefold range. Surprisingly, at the lowest concentration, cells exhibited proteomic signatures of iron shortage despite equal iron levels across conditions. We observed the uptake of labeled iron-histidine and iron-cysteine complexes, indicating that amino acids facilitated iron acquisition and that amino-acid-bound iron is bioavailable to *E. coli*. These findings reveal a previously unknown mechanism of bacterial iron acquisition that emerged under the flow imposed by the MCCD, which likely diluted the siderophore pool and reduced their efficacy. This work highlights the importance of studying bacterial physiology under low nutrient concentrations and demonstrates how physical conditions, such as flow, shape microbial nutrient acquisition strategies.

## Main Text

Bacteria constitute one of the largest biomass fractions on the planet (*1*), contribute to host health and disease (*2*), and play essential roles in biogeochemical cycles (*3*). In environments ranging from the gut to the ocean, bacteria often encounter low concentrations of diverse nutrients that limit their growth (*4*–*9*). Understanding bacterial growth under nutrient-limited conditions is challenging for multiple reasons. First, mimicking such conditions in the laboratory for extended periods is difficult: although chemostats are commonly used to study low-nutrient regimes, the use of low dilution rates can lead to inhomogeneities in nutrient distribution (*10*). Second, the interplay between multiple nutrient shortages can trigger complex physiological responses: the uptake of one nutrient may rely on the availability of another, creating a dependency in nutrient acquisition.

Proteomics has emerged as a valuable approach for mechanistically understanding bacterial physiological responses to environmental change (*11*–*15*) and diagnosing resource shortages, as one can infer underlying stressors from observed protein abundances (*16*). However, proteomics studies to date have studied a narrow range of growth conditions. While microfluidic approaches have enabled the study of bacterial physiology under low nutrient conditions (*17*), these methods typically yield insufficient biomass for proteomic analysis. Consequently, studies on bacterial physiology have primarily relied on bulk cultures with saturating nutrient concentrations and have mainly investigated how different nutrient types affect bacterial growth rates and physiological states (*12, 18*–*25*). To address this limitation, we developed a novel cultivation method that allows proteomic investigation of bacteria grown under a broad range of nutrient concentrations.

Here, we introduce the Millifluidic Continuous Culture Device (MCCD), which enables the cultivation of bacteria under constant nutrient conditions. Using the MCCD, we analyzed the proteome of *Escherichia coli* grown under different concentrations of a nutrient mixture containing amino acids, nucleobases, and vitamins. The medium provided three of the four major nutrient elements—nitrogen, phosphorus, and sulfur—in sufficient abundance to support fast growth. Carbon was supplied exclusively by the amino acids in the nutrient mixture. Our analysis revealed broad proteomic shifts across nutrient concentrations, including a reduction in amino acid biosynthesis enzymes at higher nutrient concentrations. This finding contrasts with results from studies conducted under saturating conditions that varied only the organic carbon source.

Unexpectedly, despite varying only the concentration of the nutrient mixture to impose a carbon limitation, cells in the low nutrient condition showed a proteomic signature of iron shortage. Based on these findings and the results of a chemical equilibrium model, we propose that amino acids form complexes with iron that are bioavailable—accessible for uptake and utilization—to *E. coli*. According to this hypothesis, the low abundance of amino acids in the low nutrient concentration resulted in iron limitation. This hypothesis was confirmed by uptake experiments with ^57^Fe, demonstrating that *E. coli* can acquire iron when it is complexed with cysteine or histidine. This finding thus reveals a previously unrecognized role for amino acids in facilitating iron uptake. Given the pervasive scarcity of iron across microbial habitats, our findings suggest that the role of free amino acids in iron uptake may be pervasive in microbial ecology.

### A novel device to cultivate cells for proteomics at low nutrient concentrations

We developed the Millifluidic Continuous Culture Device (MCCD) to enable the cultivation of bacteria under approximately constant nutrient conditions, even at low concentrations, by maintaining continuous fluid flow. The MCCD supports the accumulation of sufficient bacterial biomass for proteomic analysis, providing a new approach to study bacterial physiological responses to nutrient stress. Additionally, the presence of continuous flow mimics the dynamic conditions encountered in many natural microbial habitats. The device delivers a steady supply of fresh medium to bacterial cells growing within a Sterivex filter (Fig. 1A). The 0.45-µm pore size filter contained within a 2.5-mL cylinder (Fig. 1A, dotted inset) permits the flow of nutrients (2 mL/min) while retaining cells, facilitating biomass accumulation for downstream analyses.

**Figure 1.**
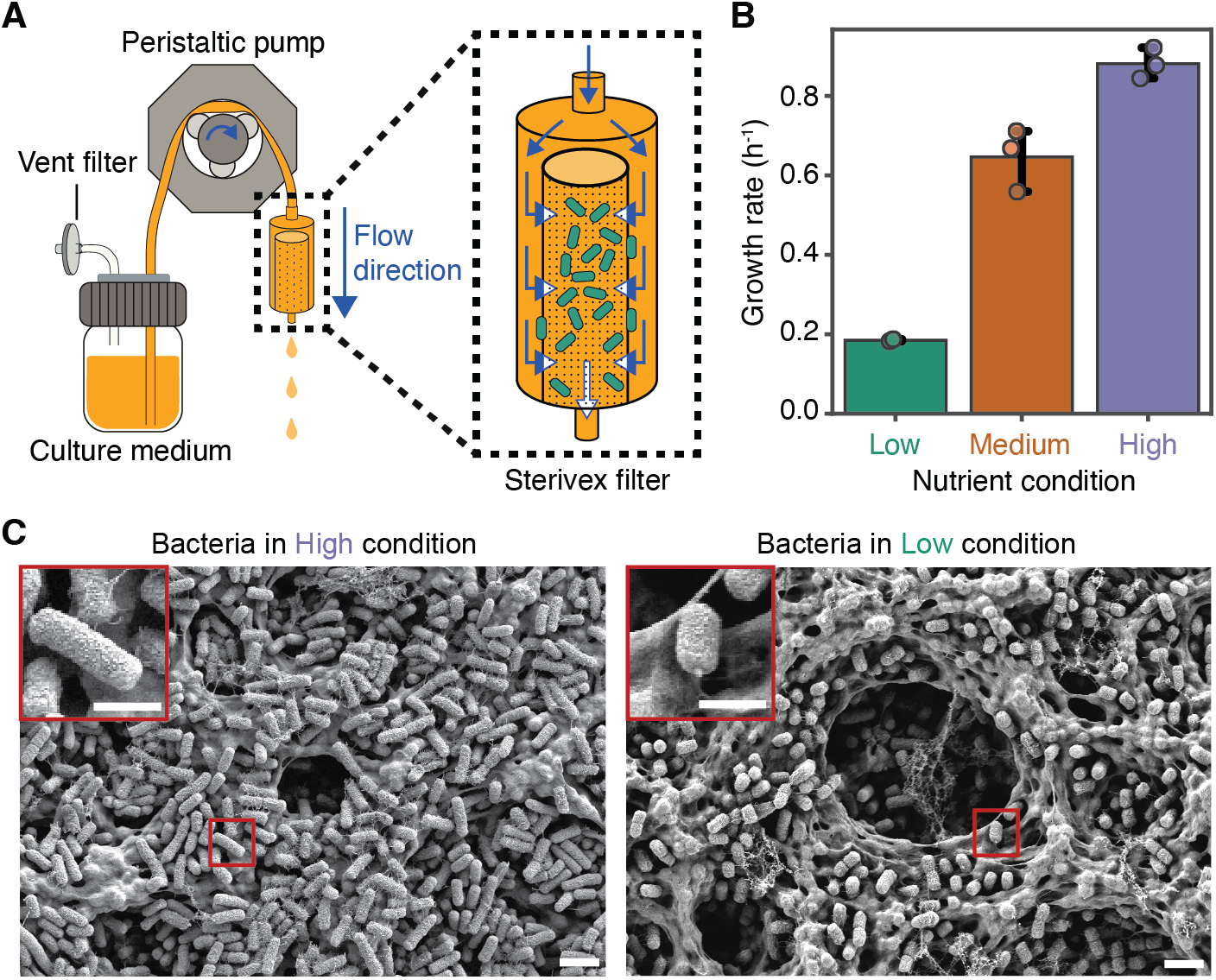
The Millifluidic Continuous Culture Device (MCCD) enables bacterial growth at controlled nutrient concentrations with sufficient biomass accumulation for proteomic analysis. (**A**) Schematic of the MCCD, with a magnified view of the Sterivex filter in which *E. coli* (green) are cultured. A peristaltic pump pushes sterile fresh culture medium at a set flow rate into the Sterivex filter through the inlet (top). A 0.45-µm pore size membrane (dotted shape) separates the filter inlet from the outlet (bottom), keeping cells in the filter while allowing culture medium to flow through. The medium flows into the filter (solid blue arrows) and traverses the membrane (white arrows with a blue outline), which restricts passage to the outlet, allowing only the medium to pass through while retaining bacteria, since these are larger than the pore size. (**B**) Specific growth rate of *E. coli* based on cell counts cultivated in the MCCD under Low (*n* = 2), Medium (*n* = 3), and High (*n* = 3) concentrations of rich-defined medium (RDM). Each point represents one biological replicate (*n*). The bar height represents the mean and error bars indicate the 95% confidence interval. The Low, Medium, and High conditions are represented by the colors green, orange, and purple, respectively. (**C**) Scanning electron microscopy (SEM) images of *E. coli* after 5 and 12 hours of growth in the High and Low conditions, respectively. The red insets show magnified views of representative bacteria. Scale bars: 2 µm (main view) and 1 µm (magnified view).

We grew *E. coli* in a MOPS-based medium (*26*) with three concentrations of a rich-defined medium (RDM) supplement to support distinct physiological states. The supplement, which provided 20 amino acids as the sole carbon source, four nucleobases, and vitamins, was diluted to maintain constant relative concentration ratios across conditions (table S1). The “High” condition contained the highest supplement concentration, the “Medium” condition contained the supplement at a 2-fold dilution relative to the High, and the “Low” condition contained the supplement at a 50-fold dilution relative to the High. Using the MCCD, we observed exponential growth in all conditions (fig. S1), with growth rates of 0.18 ± 0.002 h^−1^, 0.64 ± 0.06 h^−1^, and 0.88

± 0.03 h^−1^ in the Low, Medium, and High conditions, respectively (Fig. 1B). The MCCD sustained exponential growth for at least three doublings in each condition. Since protein half-lives are governed by cell division (*27*), we assumed that this timescale was sufficient for the proteome to adjust to the nutrient environment.

We performed experiments and simulations to characterize the nutrient environment experienced by cells in the MCCD. Scanning electron microscopy (SEM) of the bacteria on the filter confirmed that cells were distributed across the membrane surface in layers that were three cells thick or less when the culture inside the Sterivex reached approximately 10^9^ cells/mL (Fig. 1C, fig. S2). This spatial distribution ensures minimal nutrient concentration decrease between the bottom and top layers of cells (Supplementary Text 1). Numerical simulations of the flow field in the filter revealed uniform flow velocity across the membrane surface (fig. S3A) and indicated that the pressure experienced by the bacteria due to flow of growth medium was small (6.4 kPa; fig. S3B) (*28*). Finally, metabolomic analysis of the medium flowing in and out of the filters showed that, for the nutrients detectable by mass spectrometry, bacteria consumed up to 40% of some nutrients in the High condition (fig. S4). Complete nutrient exhaustion was not observed. In the Medium and Low conditions, nutrient concentrations were below the detection limit at inflow, preventing an assessment of their consumption. Together, these data indicate that the MCCD is suitable for exposing bacteria to approximately homogeneous and constant nutrient conditions.

### Evidence of a non-carbon nutrient shortage in the lowest nutrient condition

To determine the relative protein abundances in *E. coli* cells cultured in one of the three nutrient concentrations, we performed liquid chromatography tandem mass spectrometry on each of four replicate samples for each concentration. From the mass spectrometry data, we identified 1,441 proteins that account for 32% of the predicted protein-encoding genes in *E. coli* NCM3722 (Data S1, table S2). We report protein abundances as normalized spectral counts to account for variation in total spectral counts across samples, thus allowing comparison of relative protein abundances between samples. To ensure our analysis was robust to biological noise, we focused further analysis on a subset of 861 proteins that exceeded an abundance threshold (Materials and Methods). We report the abundances of these 861 proteins as percentages of total spectral counts throughout the figures and text.

To identify the biological processes with higher or lower protein abundances in each nutrient concentration, we used the EcoCyc database (*29*) to group the 861 proteins into functional protein groups (table S3). This analysis identified 52 functional groups, with 33 showing significant differences between nutrient concentrations (*p* < 0.05, Kruskal-Wallis test) (Fig. 2, fig. S5). We focused on these 33 functional protein groups (representing 627 proteins) for the remainder of the study. We conducted a comparative analysis of the Low and Medium conditions relative to the High condition, which was the least limiting for bacterial growth.

**Figure 2.**
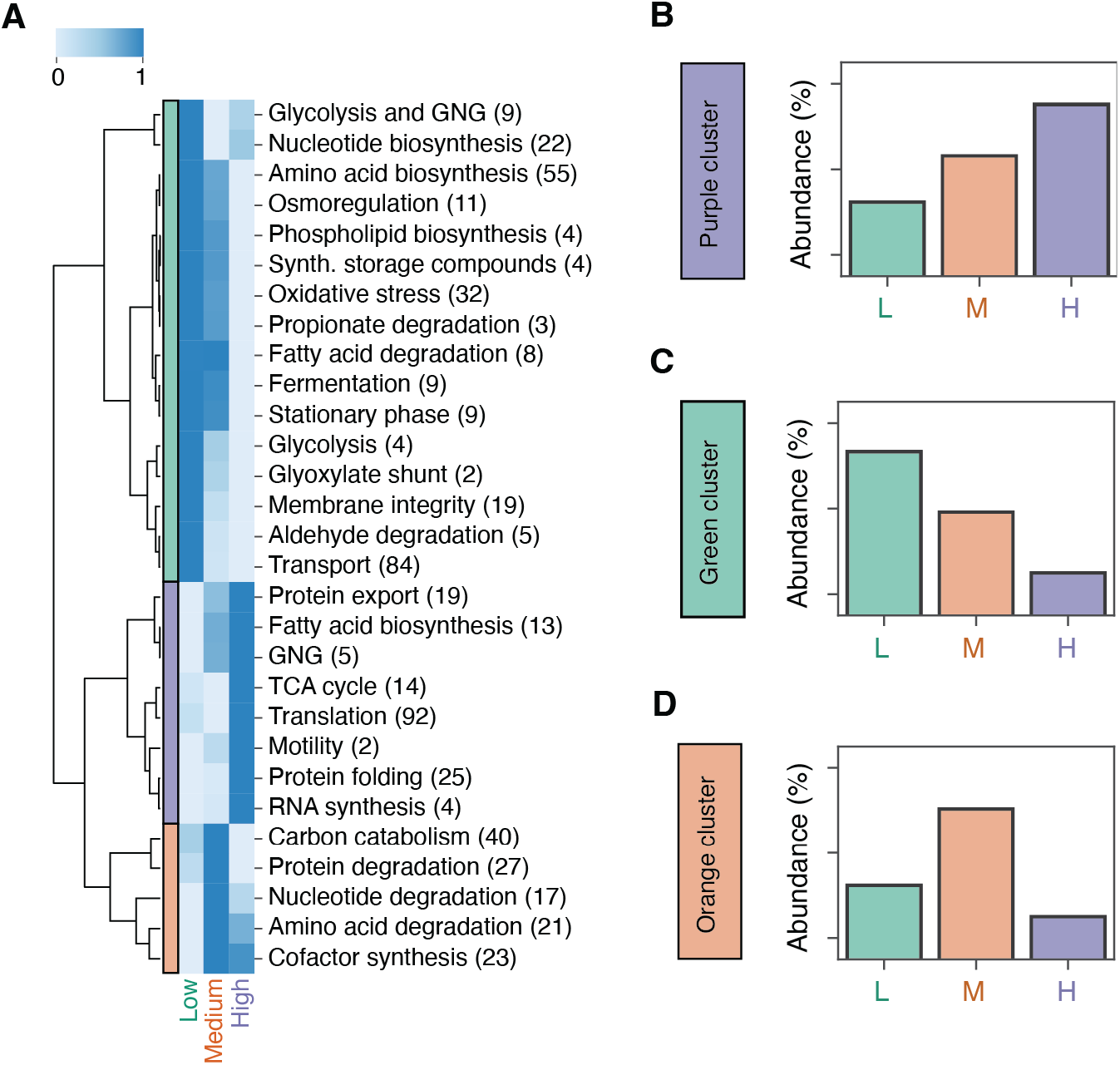
Protein expression profiles of functional groups under the three nutrient conditions. After hierarchical clustering, three groups of protein expression profiles emerged, depending on whether the highest protein abundance occurred in the Low (green bar), Medium (orange bar), or High (purple bar) nutrient conditions. Each row in the blue-shaded heatmap represents a biological function, encompassing proteins that contribute to that function. For each row, the following steps were applied: (1) the abundances of all proteins in the biological function group were summed for each nutrient condition, (2) the median value across four biological replicates within each nutrient condition was calculated, and (3) the median values were scaled across the row to range between 0 (lightest blue, lowest value) and 1 (darkest blue, highest value). Protein groups included in the heatmap met two criteria: they showed statistically significant differences across conditions (p < 0.05, Kruskal-Wallis test) and contained at least two proteins. Numbers in parentheses indicate the number of proteins in each group. See Fig. S5 for the complete heatmap. (**B-D**) Schematic bar plots illustrating the three general trends in expression profiles for each cluster that emerged in the heatmap. These representations depict how the summed protein abundances for a given functional group vary across the Low, Medium, and High nutrient conditions in each cluster.

Hierarchical clustering of the functional protein groups revealed distinct abundance patterns across nutrient concentrations (Fig. 2). From these patterns, three main clusters emerged: protein groups that increased in abundance with higher nutrient concentration (Fig. 2A,B, purple cluster), those that decreased (Fig. 2A,C, green cluster), and those that peaked in the Medium concentration (Fig. 2A,D, orange cluster). Most protein groups that increased in abundance with higher nutrient concentration (Fig. 2B, purple cluster) are associated with the demands of rapidly growing cells, such as translation, protein folding, and fatty acid biosynthesis. This cluster also included gluconeogenesis and TCA cycle enzymes, reflecting the use of amino acids as the carbon source in our experiments (*14, 30*). *E. coli* catabolize amino acids to pyruvate and TCA cycle intermediates, which feed into the TCA cycle and gluconeogenesis pathways (*14, 30*). Thus, the higher abundance of these enzymes indicates greater amino acid flux through these pathways.

Conversely, some groups that increased in abundance with lower nutrient concentration (green cluster in Fig. 2A,B) are part of the general stress response triggered by nutrient limitation (*31*). These responses protect cells from stresses and include osmoregulation, oxidative stress, and membrane integrity. Proteins in the σS regulon are known to become more abundant at slower growth rates (*22, 31*). Our observation of increased stress response proteins in the Low and Medium conditions aligns with this, confirming that the modulation of σS activity is consistent with the slower growth rates observed in these conditions. We also observed an increase in amino acid and nucleotide biosynthesis groups with lower nutrient concentrations (Figure 2A). This observation contrasts with studies of *E. coli*’s growth rate with saturating concentrations of different carbon sources or varying the uptake flux of a fixed carbon source, where amino acid and nucleotide biosynthesis enzymes are positively correlated with growth rate (*22, 25, 32*). The opposite trend in our study, where we varied the concentration of the nutrient mixture, likely reflects a decreased biosynthetic requirement for amino acids and nucleotides in the Medium and High conditions, as these are provided in the culture medium.

Unexpectedly, we observed a non-monotonic behavior in the carbon catabolism, nucleotide degradation, and protein degradation protein groups (orange cluster in Fig. 2A,D; Fig. 3A-C), with protein abundance peaking in the Medium nutrient condition. We also observed this expression pattern in more than 50% of the individual proteins in these groups (Fig. 3D-F fig. S6; Materials and Methods and table S4). An increased abundance of these protein groups is indicative of energy limitation (*31, 33*), which in our experiments is likely provided through the carbon in the amino acids. Hence, the non-monotonic expression in these protein groups indicates that cells in the Low condition experienced a shortage of a nutrient other than carbon.

**Figure 3.**
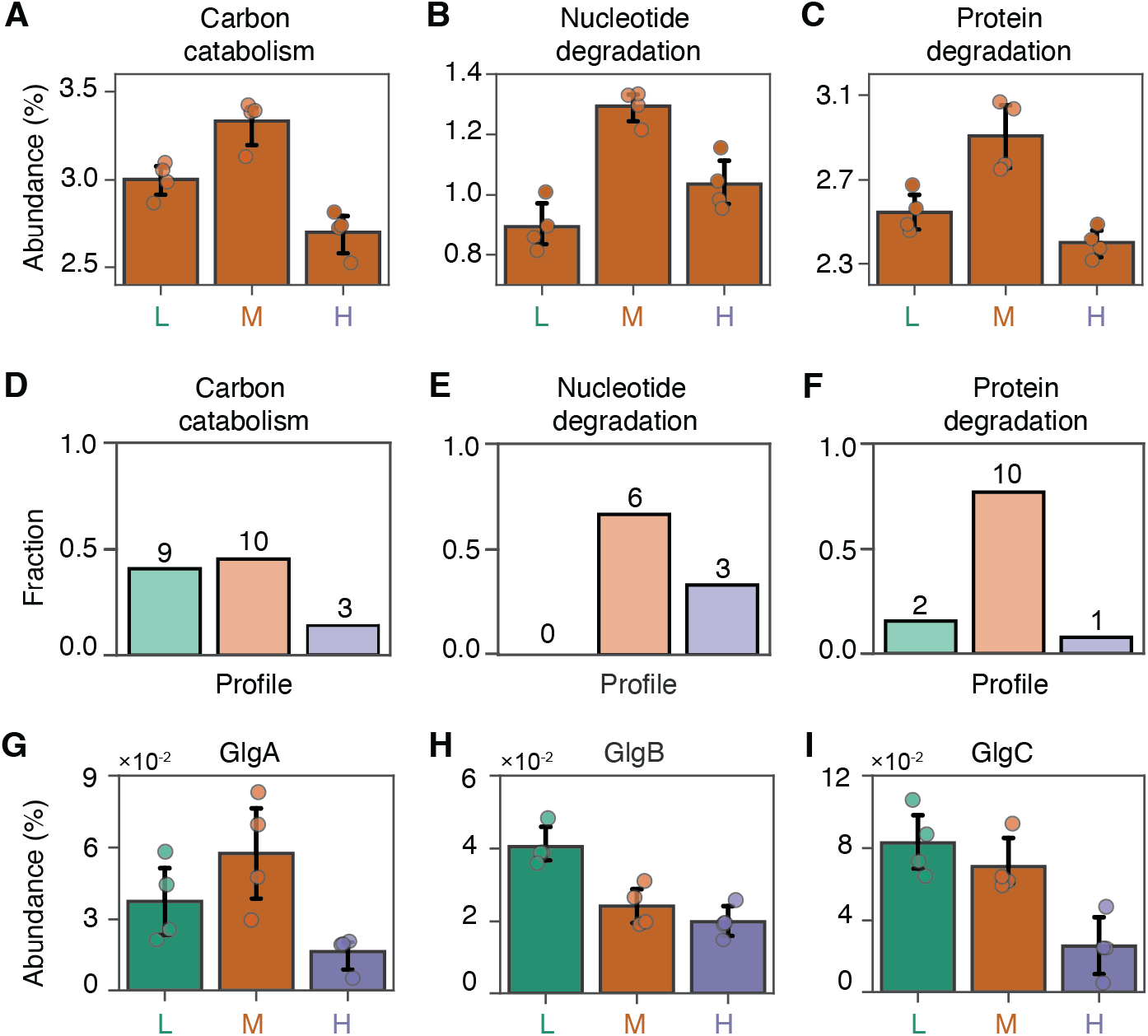
Protein expression suggests that carbon limits growth in the Medium condition but not in the Low condition. (**A-C**) Summed abundances of (**A**) all proteins assigned to carbon catabolism (40 proteins), (**B**) nucleotide degradation (17 proteins) and **(C)** protein degradation (27 proteins). For each functional group, protein abundances were summed, including proteins with and without statistically significant changes across conditions. The bar height denotes mean value, circles represent biological replicates, and error bars indicate the 95% confidence interval. Abundance is shown as the percentage of all summed spectral counts. (**D-F**) Fraction of proteins within each functional group assigned to the protein expression profiles identified in Fig. 2: highest abundance in Low (pastel green), highest abundance in Medium (pastel orange), and highest abundance in High (pastel purple). The bar height represents the fraction of proteins from that functional group displaying each expression profile, with the number above each bar indicating the number of proteins displaying that profile. Only proteins with significant differences across conditions (p < 0.05, Kruskal-Wallis test) were included (see Materials and Methods and fig. S6 for classification method). (**G-I**) Abundances of glycogen biosynthesis enzymes across nutrient conditions. The bar height denotes mean value, circles represent biological replicates, and error bars indicate the 95% confidence interval. (**A-C, G-I**) Data from Low, Medium, and High nutrient conditions are represented by green, orange, and purple colors, respectively.

Interestingly, we observed the highest abundance of enzymes involved in glycogen biosynthesis in the Low condition (“Synthesis of storage compounds” in the green cluster in Fig. 2 and Fig. 3G-I). *E. coli* synthesize glycogen when there is sufficient carbon but growth is constrained by the lack of another essential nutrient (*5, 34, 35*). The increased abundance of two key enzymes in the glycogen biosynthesis pathway in the Low condition (Fig. 3H,I), especially GlgC – the first rate-limiting enzyme – suggests that, while cells in the Low condition were the most nutrient-limited, growth was not limited by carbon. Together, the proteomic evidence indicates that nutrient limitation in the Medium condition presents as carbon limitation relative to the High condition, and as a non-carbon nutrient limitation in the Low condition relative to the Medium condition.

### Evidence for iron shortage in the Low condition

Our analysis revealed three lines of evidence of iron scarcity in the Low condition. This evidence is based on the expression patterns of the iron acquisition protein FepA, Isc system proteins, and TCA cycle enzymes (Fig. 4A). First, the iron acquisition protein FepA was more abundant in the Low condition compared to the Medium (4.4 fold) condition and the High condition (3.6 fold) (Fig. 4B). FepA is an outer membrane transporter induced under low-iron conditions, responsible for transporting enterobactin—a siderophore secreted by the cell to capture ferric iron with high affinity—into the periplasm (*36, 37*). Given that the ferric uptake regulator (Fur) represses *fepA* transcription when iron concentrations are sufficient (*36, 37*), the elevated abundance of FepA indicates iron availability was insufficient in the Low condition, triggering upregulation of this iron acquisition pathway.

**Figure 4.**
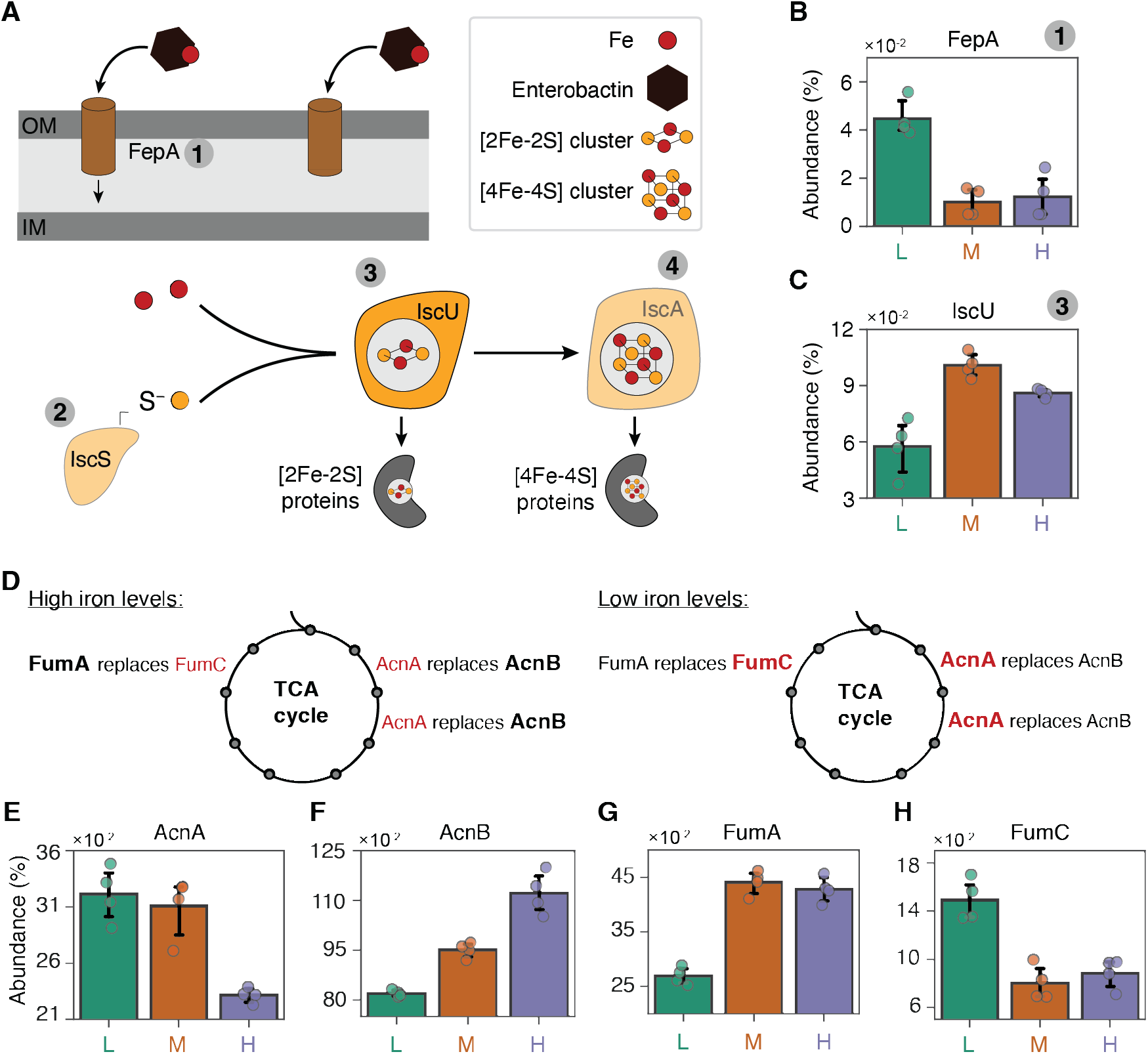
Evidence that iron, not carbon, was the limiting nutrient in the Low condition. (**A**) Schematic of the ferric enterobactin uptake system in *E. coli* and the use of intracellular iron by the Isc system. The transporter FepA (brown outer membrane protein) binds Fe(III)-enterobactin complexes and transports them into the periplasm. In the cytoplasm, iron is used by the Isc system to assemble iron-sulfur clusters (yellow and red networks), which are essential for the catalytic activity of client proteins receiving the clusters. Iron bound to enterobactin is Fe(III), while all other red dots symbols of Fe can be Fe(II) or Fe(III). (**B-C**) Barplots of the protein abundances of the ferric enterobactin transporter and the IscU system protein IscU across nutrient conditions. (**D**) Schematic of TCA cycle enzymes in *E. coli*, highlighting aconitases (AcnA, AcnB) and fumarases (FumA, FumC), which vary in protein abundance with intracellular iron levels. Enzymes in black lose catalytic activity when iron is low, while their isozymes in red preserve catalytic activity in low iron. Bold letters indicate the enzymes whose abundance increases in response to the specific iron condition (underlined). (**E-H**) Barplots of the protein abundances of the four TCA cycle iron-dependent enzymes across nutrient conditions. All barplots show individual protein abundances for each replicate (circles), along with the mean (bar height) and the 95% confidence interval (error bars). All the proteins shown in this figure displayed significant differences across conditions (p < 0.05, Kruskal-Wallis test).

Second, the expression patterns of the Isc system proteins—a multiprotein complex that assembles iron-sulfur clusters and transfers them to proteins for catalytic function and structural stability (*38*)—further support iron scarcity in the Low condition. We found that IscU, a component of the Isc system, was least abundant in the Low condition, with a 35% decrease relative to the High condition (Fig. 4C). IscS showed a similar trend, though this was not strictly significant (Kruskal-Wallis test, *p* = 0.063, fig. S7). These observations are consistent with iron scarcity, as the Isc system is known to be repressed under iron scarcity (*39*). In *E. coli*, downregulation of Isc proteins is part of the iron-sparing response, which conserves iron for critical functions by reducing the expression of iron-rich proteins when iron is scarce (*40*).

Third, in the Low condition we observed increased abundances of certain TCA cycle enzymes known to retain activity under iron scarcity (*41*–*43*). Fumarase A (FumA) and Fumarase C (FumC), which catalyze the same reaction, showed opposing expression patterns: FumA was least abundant in the Low condition, whereas FumC was most abundant (Fig. 4G,H). A similar trend was observed with the aconitase isozymes: Aconitase B (AcnB) was most abundant in the High condition, followed by the Medium condition, whereas Aconitase A (AcnA) was less abundant in the Low and Medium conditions (Fig. 4E,F). These expression patterns are consistent with the iron dependency of these enzymes. FumC remains active under iron scarcity because, unlike FumA, it lacks an iron-sulfur cluster (*41, 42*). Similarly, while both AcnA and AcnB have iron-sulfur clusters, AcnA is known to be more stable under low iron conditions, whereas the exposed cluster in AcnB is prone to dissociation (*43*). The concurrent upregulation of FumC and AcnA, alongside the downregulation of FumA and AcnB, reflects a signature iron-sparing response in *E. coli* (Fig. 4D) (*39, 44*). Together, the simplest explanation for the proteome profile in the Low condition is that the cells in the Low condition experienced iron scarcity.

### Iron-amino-acid complexes as bioavailable iron sources

The scarcity of iron in the Low condition is surprising, given that the same amount of iron (0.01 mM iron sulfate) and tricine (4 mM) was present in all nutrient conditions (Materials and Methods, (*26*)). We hypothesized that amino acid concentrations influenced iron availability due to the formation of iron-amino-acid complexes. The solubility of iron is strongly influenced by its redox state. Ferrous iron (Fe(II)) is soluble, but in oxic environments it rapidly oxidizes to ferric iron (Fe(III)), which precipitates as nearly insoluble hydroxides (occurring at concentrations around 0.5 nM in natural waters (*45*)). However, iron can form complexes with organic compounds, including certain amino acids (*46*–*50*), which, when bound as Fe(II), prevent its oxidation and maintain a soluble Fe(II) pool. In the medium we used tricine, which serves to prevent the oxidation of Fe(II) and keep it in solution (*26*). However, while tricine concentrations were constant across conditions, amino acid concentrations varied. Thus, differences in the pool of available Fe(II) complexes were likely driven by variations in amino acid concentrations.

To investigate the speciation of Fe(II), we used MINEQL+ 5.0 (*51, 52*) to construct a chemical equilibrium model and calculate the concentrations of Fe(II) complexes in each condition. Specifically, we examined two fractions of the ferrous iron pool: Fe(II)’ and Fe(II)L. Fe(II)’ represents the Fe(II) bound to inorganic compounds and the divalent cation species, Fe^2+^. Fe(II)L represents Fe(II) bound to organic ligands, including iron-amino-acid complexes. The model used the concentrations of 19 amino acids (asparagine was excluded due to the lack of stability constants), tricine, and inorganic salts for each nutrient condition, based on the defined composition of the medium. Stability constants for amino acids were obtained from the MINEQL+ database (*47, 48*). To our knowledge, stability constants for tricine have not been reported. Therefore, we substituted bicine in the model due to its structural similarity to tricine. A ligand exchange experiment confirmed comparable stability constants for bicine and tricine (Fig. S8), supporting the validity of this substitution. We set the total amount of iron as Fe(II), assuming minimal oxidation to Fe(III) based on oxidation rate estimates (Supplementary Text 2). Our results showed that the concentrations of Fe(II)-bicine ––95.8–98.7% of Fe(II) pool––and Fe(II)’ (Fe^2+^ constituting approximately 98% of Fe(II)’; supplementary table S6) remained nearly constant across the Low, Medium, and High conditions (Fig. 5A). This near-constancy indicates that tricine-bound Fe(II) and inorganic Fe(II) species were not responsible for the reduced iron availability observed in the Low condition.

**Figure 5.**
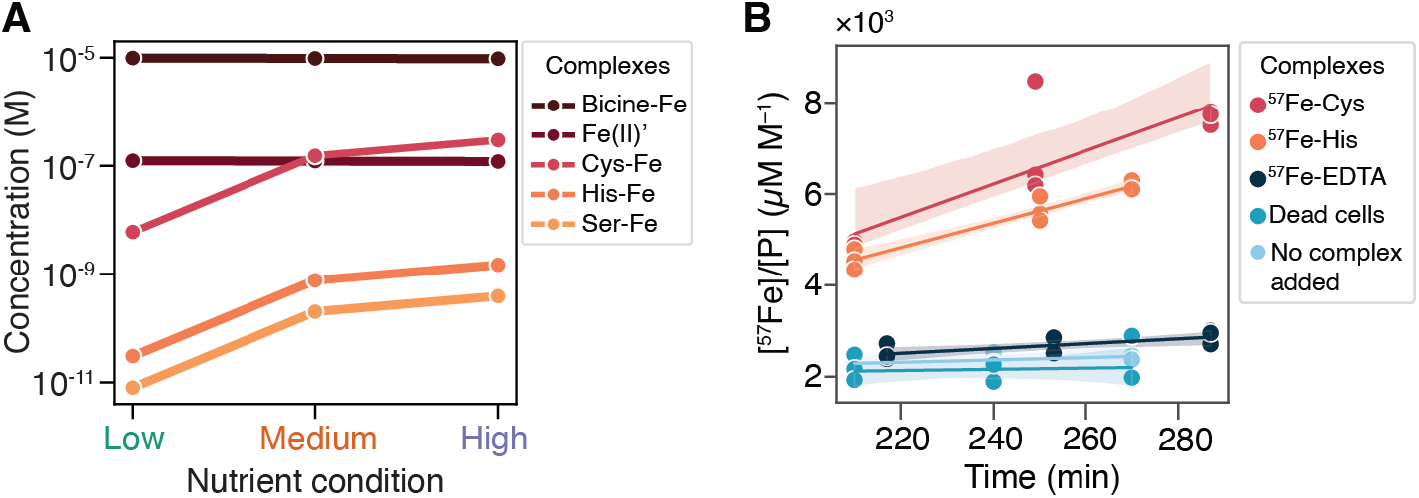
Iron uptake by *E. coli* via complexation with cysteine or histidine. (**A**) Fe(II) modeled in MINEQL yielded the concentrations of complexes between Fe(II) and components in the growth medium. Only the three highest concentration amino-acid-Fe(II) complexes are shown. Fe(II)’ represents the sum of inorganic species including Fe^2+^. (**B**) Time course of iron uptake, where tracer ^57^Fe in biomass was normalized to phosphorus ([^57^Fe]/[P]). ^57^Fe was added as complexes with cysteine, histidine, or EDTA (low bioavailability control). Glutaraldehyde-killed cells were used as a “dead cells” control, to which the ^57^Fe-Cys complex was added. Samples were collected at three timepoints over 90 minutes, with the first complex added at 210 min after growth initiation in the MCCD. Linear regression analysis yielded slopes of 36 and 27 µM M^-1^min^-1^ for the ^57^Fe-Cys and ^57^Fe-His treatments, respectively. For the controls, the slopes were 5 and 1 µM M^-1^min^-1^ for the ^57^Fe-EDTA and dead cells controls, respectively. The shaded region indicates the 95% confidence interval of the regression estimate.

In contrast, the model predicted that the low amino acid concentrations in the Low condition resulted in substantially reduced levels of Fe(II) complexed with amino acids compared to the Medium and High conditions (Fig. 5A). The concentration of the Fe(II)-cysteine complex, the most abundant Fe(II)-amino-acid complex, was two orders of magnitude lower in the Low condition (0.006 µM) than in the High condition (0.297 µM). Similarly, Fe(II)-histidine and Fe(II)-serine, the next most abundant complexes, also showed concentrations two orders of magnitude lower in the Low condition compared to the High condition (Fig. 5A, table S6). Though Fe(II)’ is considered to be a highly bioavailable form of iron (*53*), our proteomic and modeling data indicate that the iron shortage arose from the lower concentrations of iron-amino-acid complexes in the Low condition. Overall, our modeling results suggest that iron-amino-acid complexes serve as important bioavailable iron sources for *E. coli* and their scarcity in the Low condition led to iron scarcity.

### Cysteine- and histidine-iron complexes are bioavailable iron sources for *E. coli*

To test whether iron-amino-acid complexes serve as bioavailable iron sources for *E. coli*, we conducted iron uptake experiments using the rare ^57^Fe isotope, which constitutes only 2% of naturally occurring iron. Complexes of ^57^Fe-Cysteine (^57^Fe-Cys) and ^57^Fe-Histidine (^57^Fe-His) were prepared by mixing ^57^Fe with cysteine or histidine at a 1:10 molar ratio and spiked into actively growing cultures in the MCCD at three timepoints, spaced 30 minutes apart. The final ^57^Fe concentration in the culture was 5 µM (compared to 10 µM of ^56^Fe in the medium), and cysteine or histidine reached 50 µM (compared to 0.5 µM and 1 µM in the medium, respectively). Controls included (1) a “dead cells” control with glutaraldehyde-killed cells spiked with the ^57^Fe -Cys complex, (2) a non-bioavailable control with an ^57^Fe -EDTA complex, and (3) no iron complex addition. At each sampling timepoint (20 min after spiking; see fig. S9 for experimental design schematic), cell pellets were harvested, and intracellular iron (^56^Fe and ^57^Fe isotopes) concentrations were quantified using inductively coupled plasma mass spectrometry (ICP-MS).

The results showed that ^57^Fe -amino-acid complexes were bioavailable to *E. coli*. To account for biomass differences, ^57^Fe concentrations were normalized to phosphorus levels ([^57^Fe]/[P]), assuming proportionality between phosphorus content and cell growth (*18, 54*). A linear regression analysis revealed that the slopes of [^57^Fe]/[P] for the ^57^Fe -Cys and ^57^Fe -His treatments were 36 and 27 µM M^-1^min^-1^, respectively. In contrast, the ^57^Fe-EDTA and dead-cells control had slopes of 5 and 1 µM M^-1^min^-1^ (Fig. 5B). These results indicate that ^57^Fe accumulation in the ^57^Fe -Cys and ^57^Fe -His treatments was active and mediated by amino acid complexation. The unchanged [^57^Fe]/[P] levels in the dead-cell control confirmed that uptake of iron-amino-acid complexes was not passive, and the unchanged levels in the ^57^Fe-EDTA control, acting as the non-bioavailable control, demonstrated that iron uptake was not driven by non-specific chelation.

We further examined whether *E. coli* preferentially take up amino acid-bound iron. We compared the rates of iron uptake between ^57^Fe, delivered as pre-formed iron-amino-acid complexes, and the most naturally abundant isotope, ^56^Fe, predominantly bound to tricine in the medium (Fig. 5A). We interpret differences in the ratio of intracellular ^57^Fe to ^56^Fe as differences in uptake efficiency, with a higher ^57^Fe/^56^Fe ratio providing evidence of facilitated iron uptake due to complexation with amino acids. We used an ordinary least squares (OLS) linear model to analyze the time-course measurements of intracellular ^56^Fe and ^57^Fe concentrations for each treatment. The analysis revealed that the uptake rate of ^57^Fe was 3.8 times higher than that of ^56^Fe in the ^57^Fe-Cys treatment and 5.8 times higher in the ^57^Fe-His treatment (*p* < 0.05, two-tailed t-test, Fig. S9A,B). These results indicate that the complexation of iron with cysteine or histidine facilitates iron uptake.

## Discussion

Using a novel bacterial culturing device (the MCCD), we studied the physiology of *E. coli* using proteomics in environments with different nutrient concentrations in the presence of continuous flow. In the Low condition, proteomic analysis revealed an increased abundance of siderophore-mediated iron uptake proteins and changes in iron-dependent enzymes, indicating an iron shortage. A chemical equilibrium model showed that iron-amino-acid complexes varied more than other iron species, suggesting that these complexes contribute significantly to the bioavailable iron pool. The observed iron uptake from cysteine- and histidine-bound ^57^Fe in uptake experiments revealed that iron-amino-acid complexes are bioavailable to *E. coli*. Our findings demonstrate an unanticipated dependency of iron uptake on amino acid uptake.

We posit that iron-amino-acid complexes constitute a significant fraction of bioavailable iron in natural environments where siderophores are highly diluted. Although characterized decades ago (*46*–*48*), the role of these complexes in iron uptake has been overlooked, likely due to the prevalent use of batch cultures in bacterial physiology studies. In batch cultures, the confined environment allows for the accumulation of high-affinity organic chelators, such as siderophores, which mediate Fe(III) uptake by binding and transporting Fe(III) into the cell via specific receptors (fig. S10A) (*37, 55*). In contrast, Fe(II) uptake occurs via diffusion across outer membrane porins and subsequent transport into the cytoplasm via the Feo system (fig. S10B) (*56*). In contrast to batch cultures, many natural microbial habitats are unconfined or subject to fluid flow, preventing siderophore accumulation and thus hampering this iron acquisition pathway. In our study, the continuous flow in the MCCD device likely acted against siderophore accumulation, allowing us to discover a role of iron-amino-acid complexes in iron uptake.

The effectiveness of iron uptake strategies is context-dependent. Siderophores, with extremely high iron binding affinity, are likely advantageous in confined or physically structured environments such as particles, which act as structured islands that limit the washout of siderophores by flow (*57*–*60*). In contrast, in unconfined environments like the open ocean, diffusional losses make siderophore use less efficient unless the concentration of secreting cells is large. It has been estimated that a cell must secrete approximately 28,000 siderophores to capture one single iron ion (*61*). Even though this can be partially mitigated by collective siderophore secretion (*61*), it still imposes a significant nitrogen and carbon cost (*57, 61*). To reduce diffusive losses, most marine bacteria use amphiphilic siderophores –siderophores with cell membrane affinity (*62*) – or employ membrane-embedded uptake systems, such as those for Fe(II), as alternative strategies for iron acquisition. The majority of soluble iron is complexed with organic ligands, categorized as “strong” (e.g., siderophores) or “weak” (e.g., amino acids) (*63, 64*). Weak ligands, including amino acids, are abundant, biosynthetically inexpensive, and may facilitate iron uptake with lower metabolic cost relative to siderophore production. Saccharides, another class of weak ligands, enhance iron uptake in phytoplankton (*65*). Similarly, our findings suggest that amino acids, acting as weak ligands, enable iron acquisition when taken up as metabolites by bacteria, and we propose that this uptake mechanism is particularly important in unconfined environments where siderophores are less effective.

Iron-amino-acid complexes could be taken up either through specific transporters for these complexes or via amino acid transporters that might also allow iron hitchhiking on amino acids. Our proteomic data showed that the *E. coli* outer membrane porins OmpF and OmpC, which facilitate the transport of small molecules such as amino acids and Fe(II) into the periplasm (*56*), were most abundant in the Low condition (fig. S11). However, the observed changes in transporter abundance in our study likely resulted from the combined effects of reduced iron and amino acid availability. While specific iron-amino-acid uptake systems in bacteria have not been identified, similar transport systems have been described for other metals: both *E. coli* and *Staphylococcus aureus* possess ABC transporters that mediate the uptake of histidine-bound nickel (*66, 67*). Future work may reveal if iron-amino-acid complexes uptake occurs through previously characterized amino acid transporters or through dedicated transporters.

We propose that the iron uptake mechanism described here plays a significant role in environments where fluid flow or diffusion limits siderophore accumulation. In humans, the gut experiences fluid flow rates of up to 20 mL/min in the small intestine (*68*) and 1.5 mL/min in the colon (*69*), potentially hindering the accumulation and effectiveness of siderophores. Iron is a limiting nutrient in the gut because the majority is sequestered by high-affinity host proteins, leaving only trace amounts available to microbes (*70*). At the same time, dietary protein digestion releases peptides and free amino acids, providing a carbon and nitrogen source for gut bacteria like *E. coli* (*6*). By coupling the uptake of amino acids with iron bound to them, bacteria may acquire iron without the metabolic expense of siderophore production. Siderophores require substantial carbon and nitrogen for biosynthesis (*71*), and their loss to flow and diffusion further amplifies their cost. In contrast, the uptake of iron through iron-amino-acid complexes represents a more economical strategy for microbial iron acquisition in such nutrient-limited, dynamic environments.

The uptake of iron via iron-amino-acid complexes is likely widespread among bacteria, as transport systems like ABC transporters are highly conserved across bacterial species (*72*). In marine environments iron is a limiting nutrient that constrains primary productivity and bacterial heterotrophs (*73*). Amino acids support 5–55% of bacterial nitrogen and carbon demands in these environments (*74*–*77*). The SAR11 clade, for instance, assimilates up to 60% of available amino acids in oligotrophic waters (*77*), supported by high-affinity transport systems (*78*). Given the important role of amino acids as carbon and nitrogen sources, the iron uptake mechanism described here is likely broadly relevant, enabling bacteria to acquire iron through amino acid transport.

Our study reveals a striking example of iron uptake dependency on amino acid availability influenced by environmental physical conditions: as fluid flow dilutes the siderophore pool, amino acids become essential iron ligands for uptake, triggering iron scarcity in iron bioavailability under low amino acid concentrations. In the absence of flow, siderophore accumulation would likely mitigate this dependency. By replicating key features of natural microbial habitats—low nutrient concentrations and fluid flow—we uncovered a previously unknown mechanism of bacterial iron uptake and demonstrated how physical conditions influence nutrient acquisition strategies.

## Acknowledgments

We thank Martin Ackermann, Julia Vorholt, the current and former members of the Stocker group, and members of the PriME collaboration for insightful discussions. We are grateful to Suckjoon Jun for generously providing the NCM3722 strain of *E. coli* used in this study. We thank Miriam Lucas from ScopeM for processing samples for SEM and acquiring the SEM images, as well as the entire ScopeM team for their support and technical assistance. We thank Dawn Moran for processing samples for proteomic analysis. We also thank Samuel Charlton and Eleonora Secchi for their assistance in planning and executing the Sterivex flow calibration experiments and Andrés Velásquez-Parra for assistance in visualizing flow velocity and pressure from the numerical model. Finally, we thank Russell Naisbit for his critical review, which significantly improved this manuscript, and we acknowledge the use of ChatGPT-4 for grammar correction and phrasing enhancements.

## Author contributions

Conceptualization: JLG, RS, MS, JN

Methodology: JLG, RS, MS, JN, UA, ZL, JJM

Investigation: JLG, MRM, IS, DMM, SP, JJM, JN, UA, ZL, JMK

Visualization: JLG, JMK, JN, IS, JJM

Funding acquisition: RS, MS, TH, US

Formal analysis: JLG, MRM, IS

Supervision: RS, MS

Writing – original draft: JLG, JN, JMK

Writing – review & editing: JLG, RS, MS, JN, JMK, UA, TH

## Data and materials availability

All data and code will be available upon publication.

## Supplementary Materials

Materials and Methods

Supplementary Text S1 to S2

Figs. S1 to S10

Tables S1 to S6

References

Data S1

