## Supplementary material for "Growth in Low Carbon Conditions Reveals Amino-Acid-Coupled Iron Uptake"

**The PDF file includes:**

Materials and Methods
Supplementary Text 1
Supplementary Text 2
Figs. S1 to S12
Tables S1 to S8

**Other Supplementary Materials for this manuscript include the following:**

Data S1

### Materials and Methods

Bacterial strain

Experiments were performed using *E. coli* K-12 NCM3722 Δ*motA*.

Growth media

All growth media used in this study were based on MOPS-buffered medium (*26*). The medium is composed of 40 mM MOPS and 4 mM tricine (adjusted to pH 7.4 with 10 M KOH), 0.01 mM $FeSO_4$, 9.5 mM $NH_4Cl$, 0.276 mM $K_2SO_4$, 0.5 µM $CaCl_2$, 0.525 mM $MgCl_2$, 50 mM NaCl, 1.32 mM $KH_2PO_4$. Micronutrients were added to final concentrations of $2.92 \times 10^{-7}$ mM $(NH_4)_6Mo_7O_{24} \cdot 4H_2O$, $4 \times 10^{-5}$ mM $H_3BO_3$, $3.02 \times 10^{-6}$ mM $CoCl_2$, $9.62 \times 10^{-7}$ mM $CuSO_4$, $8.08 \times 10^{-6}$ mM $MnCl_2$, and $9.74 \times 10^{-7}$ mM $ZnSO_4$.

RDM dilutions

We prepared the Rich Defined Medium (RDM) dilutions using two supplement solutions, 5x EZ (amino acid and vitamin mixture) and 10x ACGU (nucleobase mixture), both from the MOPS EZ Rich Defined Medium (Teknova, M2105), in a 2:1 ratio to the MOPS medium. The Full RDM is made by diluting the EZ and ACGU stock solutions five and ten times, respectively, in MOPS medium.

For proteomics experiments, we selected three specific dilutions, termed High, Medium, and Low conditions, corresponding to 0.5%, 0.255%, and 0.01% of the full RDM concentration, respectively. The High medium was prepared by mixing 0.1 ml of 5x EZ and 0.05 ml of 10x ACGU in 99.85 ml of MOPS-buffered medium. The Low medium was produced by diluting 2 ml of the High medium into 98 ml of MOPS-buffered medium. The Medium medium was created by mixing equal parts of the High and Low media. The dilution process ensures that while the absolute concentrations of amino acids, nucleobases, and vitamins change across conditions, their relative concentrations remain constant. The exact concentrations of amino acids, nucleobases, and vitamins in these three media are detailed in table S1.

Seed culture preparation

To prepare the overnight (ON) cultures, we inoculated a -80 ˚C glycerol stock into 3 ml of MOPS-buffered medium supplemented to a 10% RDM concentration (60 µL 5x EZ and 30 µL 10x ACGU) and incubated at 37 ˚C and 200 rpm for 16–18 h. This ON culture was used to prepare the seed culture by diluting at a ratio of 1:1000 in the same medium. Once the seed culture reached an $OD_{600}$ of 0.07–0.1, it was used to initiate experiments.

Millifluidic Continuous Culture Device

The Millifluidic Continuous Culture Device (MCCD) employs a Sterivex filter (Millipore) as a culture vessel for bacteria. The Sterivex filter comprises a cylindrical plastic casing enclosing a porous membrane securely positioned along the inner surface of an inner plastic cylinder. An inlet port located at one end of the casing enables the loading of liquids, which traverse the membrane under applied pressure, accessing the inner cylinder. These liquids exit through the outlet port, while particles larger than the membrane's pore size are retained within the chamber between the casing and the membrane. In this study, we employed the space between the membrane and the casing as a cultivation chamber for bacteria. Sterivex filters with a 0.45 µm pore size and a biologically compatible polyvinylidene fluoride (PVDF) membrane (Millipore Catalog # SVHVL10RC) were used. A peristaltic pump (Shenchen Lab V1) continuously delivered fresh medium from a flask aerated aseptically with an vent filter (Millipore Millex-FG SLFG05000) into the Sterivex filter. For each experiment, 2 ml of seed culture were introduced into the Sterivex filter. Prior to connecting the peristaltic pump tubing, one check valve (Masterflex #MFLX30505-91) was connected to the inlet and another to the outlet of the Sterivex filter to introduce flow resistance and maintain the Sterivex in a filled state. The peristaltic pump was set to a flow rate of 2 ml/min. The flow of fresh medium through the Sterivex filter was sustained uninterrupted throughout the experiment, with outflow collected in a waste receptacle.

The collection of samples was performed by unplugging the Sterivex filter from the tubing and introducing it into a 50 mL Falcon tube with the inlet port facing down. The tubes containing the Sterivex filters were centrifuged for 2 min at 2500 rcf to collect the culture from the casing. The filters were cut using a sterile pipe cutter and rinsed to collect the cells accumulated in the casing walls. This sample collection procedure was used for the growth curve experiments and the collection of samples for proteomic analysis.

For the growth curve experiments, multiple Sterivex filters were seeded at the start of an experiment, each corresponding to a planned time point. At each time point, a Sterivex filter was unplugged, opened, and the culture was collected as described above. From the culture, 2-3 samples (100 µl each) were fixed with glutaraldehyde to a final concentration of 1.25%, stained with Sybr Green (Thermo Fischer S7563) for 10 minutes in the dark, and analyzed by flow cytometry. Forward scatter (FSC), side scatter (SSC), and green fluorescence were recorded at a flow rate of 25 µl/min. Bacterial cells were counted by averaging the results from gates applied using FSC vs. Fluorescence and SSC vs. Fluorescence.

Proteomic samples were collected using the same sample collection procedure, and cell counts per sample were determined via flow cytometry (table S2). For the High and Medium conditions, we sampled 5 h after initiating the experiments. For the Low condition, we sampled 15 h after initiation due to the slower growth rate. We performed four replicate experiments for each concentration and performed proteomic analysis on each replicate sample.

### Numerical simulations of flow through the Sterivex filter

We simulated fluid flow velocity and pressure using the finite element software Comsol Multiphysics®. The model was based on geometrical information provided by the filter's manufacturer and our own measurements. Steady-state velocity and pressure fields in the liquid phase were modeled using the Navier–Stokes equations for fast flow in pipes and the Brinkman equations for slow flow in porous media, such as the membrane. We assumed an incompressible fluid with constant density and dynamic viscosity. No-slip conditions were applied to the pipe walls and the porous membrane. As boundary conditions, we imposed the flow rate at the inlet and the pressure at the outlet.

To calibrate the model, we conducted two sets of pressure measurements. In the first set, we determined the membrane's permeability by measuring the pressure difference across a filter without bacteria at various flow rates (0.1–1 mL/min). The applied pressure was controlled using a microfluidic flow controller (Flow EZ, Fluigent), and the flow rate was measured using a bidirectional microfluidic flow sensor (Flow Unit, Fluigent). We recorded pressure differences at 20-second intervals over 20 minutes for each flow rate. The average pressure differences were then used to adjust the numerical model, yielding an estimated membrane permeability of approximately $10^{-15}$ $m^2$.

In the second set of experiments, we monitored pressure differences during bacterial growth within the filter at a constant flow rate of 1 mL/min. As *E. coli* grew over 14 hours, the effective permeability of the membrane decreased to approximately $10^{-16}$ $m^2$. These permeability values, with and without bacteria, were then incorporated into the model to simulate the velocity and pressure fields under both conditions.

<u>Metabolomics of inflow and outflow</u>

*E. coli* was grown under the three nutrient conditions in the MCCD at a flow rate of 2 mL/min. Inflow media samples were collected at the start of the experiments. After the bacteria had grown for the same duration as in the proteomics experiments (5 hours for the High and Medium conditions, and 15 hours for the Low condition), outflow media samples were collected. Following this initial collection, the flow rate was reduced to 0.2 mL/min, and additional outflow samples were taken.

The collected samples were diluted 10-fold and analyzed using LC-QTOFMS. The measurements were conducted on an Agilent 6520 Time of Flight Quadrupole Time of Flight Mass Spectrometer in negative and positive mode, high-resolution mode with a 0.9 Hz scan rate, and an acquisition mass range of 50-1,700 m/z. The drying gas flow rate was 10 L/min, nebulizer pressure was 30 psig, and gas temperature was 325°C. An Agilent 1100 series liquid chromatography stack was used to inject 3 µL of the sample into an Agilent EC-CN Poroshell column (2.7 µm, 50 × 2.1 mm) with an adapted salt-tolerant method to reduce ion suppression from salts in the medium (*79*). The sample injection order was randomized, and all samples were injected in single technical replicates. The column temperature was maintained at 20°C, and the flow rate was set to 350 µL/min. The mobile phase consisted of 10% Acetonitrile (CHROMASOLV), 90% mass spectrometry grade water, and 0.01% Formic acid (v/v), and was operated isocratically. For every 40 samples, the column was washed by flushing for 5 minutes with 90% acetonitrile (CHROMASOLV), 10% mass spectrometry grade water, and 0.01% formic acid (v/v), followed by a 10-minute equilibration with the isocratic buffer. Data analysis was performed using Agilent Quantitative MassHunter software, and compounds were quantified using a calibration curve of standards with known concentrations.

<u>Image acquisition by Scanning Electron Microscopy</u>

To evaluate distribution of cells on the porous membrane, we grew cells in the MCCD using the High and Low RDM nutrient conditions. After 5 h growth, cells were fixed by introducing a solution of 2.5% glutaraldehyde and 2% formaldehyde (both EM grade; Polysciences Europe GmbH, Hirschberg an der Bergstrasse, Germany) in 0.15 M sodium cacodylate buffer into the

Sterivex filter, and incubated overnight. Before further processing, the Sterivex filter was opened and the filter extracted, cut into strips of approx. 5 × 10 mm and these transferred into 0.15 M sodium cacodylate buffer. The fixative solution was stored to count the dislodged cells with flow cytometry. The samples were washed in fresh buffer three times, and then post-fixed with 1% osmium tetroxide (Polysciences). The samples were dehydrated in a series of solutions of increasing ethanol concentration (50%, 75%, 90%, 98%, 3x 100%). All dehydration steps after extraction from the Sterivex filter were performed in a BioWave Pro+ Tissue Processor (Ted Pella Inc., Redding CA, USA). The samples were then critical point dried from ethanol with liquid $CO_2$ using a critical point dryer (Autosamdri- 931, Tousimis, USA). Afterwards, the filter strips were mounted onto SEM stubs using carbon tape and vacuum sputter coated with 8 nm Pt–Pd (CCU-010, Safematic GmbH, Switzerland) before imaging them using a scanning electron microscope (Merlin, Zeiss). Images were acquired using an InLens or HE-SE2 secondary electron-detector with an acceleration voltage of 2 kV at a working distance of 5 mm.

### Proteomics

Tryptic peptides were analyzed via liquid chromatography tandem mass spectrometry (LC/MS/MS) using a Michrom Advance HPLC system with reverse phase chromatography coupled to a Thermo Scientific Q-Exactive Orbitrap mass spectrometer with a Michrom Advance CaptiveSpray source. Each sample was concentrated onto a trap column (0.2 × 10 mm ID, 5 µm particle size, 120 Å pore size, C18 Reprosil-Gold, Dr. Maisch GmbH) and rinsed with 100 µL 0.1% formic acid, 2% acetonitrile (ACN), 97.9% water before gradient elution through a reverse phase C18 column (0.1 × 150 mm ID, 3 µm particle size, 120 Å pore size, C18 Reprosil-Gold, Dr. Maisch GmbH) at a flow rate of 500 nL/min. The chromatography consisted of a nonlinear 220 min gradient from 5% to 95% buffer B, where buffer A was 0.1% formic acid in water and buffer B was 0.1% formic acid in ACN (all solvents were Fisher Optima grade). The mass spectrometer monitored MS1 scans from 380 *m*/*z* to 1280 *m*/*z* at 70K resolution. MS2 scans were performed on the top 15 ions with an isolation window of 2.0 *m*/*z* and a 15 s exclusion time.

Mass spectra were searched against the translated genome (*80*) using Proteome Discoverer's SEQUEST HT algorithm (Thermo) with a fragment tolerance of 0.02 Da and parent tolerance of 10 ppm. Identification criteria consisted of a protein threshold of 1% False Discovery Rate (FDR), peptide threshold of 0.1% FDR, and two minimum peptides when analyzed with Scaffold version 5.1.2 (Proteome Software, Inc.).

### Data Analysis

The proteomic data of each protein were subjected to a filtering process. First, a threshold of 5 counts was set as the value below which the data point was considered to be in the range of biological noise. Next, we asked how many of the four replicate measures per condition were above the threshold. If one of the three conditions had more than three replicate values above the threshold, the protein was retained for further analysis. With this thresholding method, 861 proteins were listed for analysis. Using information on the physiological role of each protein obtained from EcoCyc (4), we classified the proteins into functional groups (see table S3 for the complete list and group assignments). To test for significant differences across conditions for each functional group, we performed a Kruskal-Wallis test, setting the significance level (α) at

0.05. For multiple hypothesis testing correction, we applied the Benjamini-Hochberg procedure, controlling the FDR at 0.05.

The proteins in the Carbon catabolism, Nucleotide degradation, and Proteases groups that showed statistically significant changes were classified according to their abundance trends across conditions. Each protein was represented as a point in 3D space as follows: the median protein abundance in the Low condition represented the x-coordinate, the median protein abundance in the Medium condition represented the y-coordinate, and the median protein abundance in the High condition represented the z-coordinate. We then applied a K-means clustering algorithm, setting the number of clusters to eight. This choice was based on a visual inspection of protein abundance trends, which revealed eight distinct patterns (see fig. S6 for a depiction of each). The algorithm assigned each protein to one of the eight clusters shown in fig. S6. Proteins without statistically significant changes were categorized as No Change (NC).

Data preparation and analysis were conducted in Python version 3.11.8 using the Matplotlib 3.6.3, Seaborn 0.11.2, and SciPy 1.9.3 libraries. The method for hierarchical clustering in Fig. 2 was an unweighted pair group method with arithmetic mean (UPGMA) that used the "Manhattan" distance metric, implemented and plotted using Seaborn.

### Chemical equilibrium model

To investigate the speciation of iron in our experimental conditions, we employed MINEQL+ 5.0 (*51*, *52*), a chemical equilibrium modeling software. The inputs for the model included the concentrations of 19 amino acids (excluding asparagine), tricine, and inorganic salts present in the media. Stability constants for amino acid-iron interactions were sourced from the MINEQL+ database (table S5), with tricine approximated using bicine due to similar chemical properties (*47*, *48*). We ran the model with a fixed pH of 7.4 and enabled the ionic strength correction.

### Uptake experiments with the $^{57}$Fe isotope

To investigate the role of iron-amino acid complexes in iron uptake, we exposed actively growing *E. coli* cultures in the MCCD to complexes of the rare $^{57}$Fe isotope with specific ligands during three 10-minute time points. Three Sterivex filters were inoculated with a seed culture and incubated in a modified MOPS-based medium, where the iron sulfate concentration was lowered to 0.001 mM while maintaining the same nutrient concentrations as in the High condition. After 1.5 hours of growth, $^{57}$Fe complexes were prepared with cysteine and histidine at a molar ratio of 1:10 ($^{57}$Fe:amino acid), achieving final concentrations of 0.0005 mM $^{57}$Fe and 0.005 mM amino acid in the medium. After 3.5 hours of growth, the incoming medium into the Sterivex filters was replaced with the modified MOPS-based medium containing the prepared $^{57}$Fe-amino acid complex. This spiked medium was flowed through the Sterivex filters for 10 minutes at the same flow rate. Next, the flow was switched back to the original modified MOPS-based medium for an additional 10 minutes to wash away any residual complex. After the wash, cells from one Sterivex filter were harvested, centrifuged, and stored at −80 °C. This process was repeated every 30 minutes for two additional time points. Three controls were included: an EDTA control, in which $^{57}$Fe-EDTA was added to the medium at a final $^{57}$Fe concentration of 0.0005 mM and EDTA concentration of 0.01 mM; a "dead cells" control where glutaraldehyde was added to a final concentration of 1.5% to kill the cells; and a no-complex control without any $^{57}$Fe-ligand complex. The $^{57}$Fe-cysteine complex was used to spike the "dead cells" control.

We quantified $^{57}$Fe-bound cysteine and histidine uptake using inductively coupled plasma mass spectrometry (ICP-MS; Thermo Scientific™ iCAP-Q ICP-MS). Prior to analysis, 15 mL Basix™ polypropylene conical centrifuge tubes used for sample dilutions were cleaned with 10% hydrochloric acid (HCl; Fisher Chemical™ certified ACS Plus, 36.5–38.0% HCl) for 7 days, then rinsed three times with a pH 2 solution (Fisher Chemical™ certified ACS Plus, 36.5–38.0% HCl) to ensure trace-metal-clean conditions. The tubes were then weighed prior to use. Following the cleaning and weighing process, 0.5 mL of a solution containing 50% nitric acid (Fisher Chemical™ Optima™ 67–70% $HNO_3$) and 100 ppb In (diluted from Inorganic Ventures™ certified In standard at 1000 ppm) was added to experimental samples in 2 mL conical-bottom microcentrifuge tubes containing cell pellets while they thawed. The sample tubes were vortexed daily for ~20 seconds each to suspend the cell pellets and assist in digestion. The samples were then acidified for five days at room temperature to ensure complete leaching of metals and digestion of organic matter. Once the samples were digested, 0.1 mL of the sample solution was transferred to a 15 mL trace-metal-cleaned tube and diluted with 1 mL of ultrapure water (resistivity 18.2 MΩ·cm at 25 °C) to reach final In concentration of ~10 ppb in all samples. The trace-metal-cleaned tubes, sample solutions, and diluents were all weighed on an analytical scale to accurately determine the dilution factor and calculate the final concentrations in the samples. Prior to analysis on the ICP-MS, calibration standards were prepared using mixed metal and phosphorus (P) standards (certified mixed metal and P standards from Inorganic Ventures™; calibration concentrations of 0, 1, 5, 10, 20, and 100 ppb for metal standards, and 50, 100, 200, and 300 ppb for P standards). Blanks, process blanks, diluted samples, and calibration standards were all analyzed by ICP-MS to quantify $^{57}$Fe, $^{56}$Fe, and P concentrations.

The total Fe concentration in the stock solutions (Fe-57 Cysteine, Fe-57 Histidine, and Fe-57 EDTA) was determined using ICP-MS. Prior to analysis, approximately 1.5 mL of each stock solution was sampled into 2 mL microcentrifuge tubes. The stock solutions were stored at -80˚C and defrosted in an anoxic chamber immediately before dilution. For dilution, 50 µL of the stock solution was added to a trace metal-clean 15 mL Basix™ polypropylene conical centrifuge tubes (Fisher Scientific™ Catalog No.14-955-238) and diluted 400 times with a solution containing 10 ppb In (Inorganic Ventures™) in 5% nitric acid (Nitric Acid 67-70%, Optima™, for Ultra Trace Elemental Analysis, Fisher Chemical™; Catalog No.A467-250). Calibration curves were prepared using Certified Inorganic Ventures™ standards. The diluted stock solutions, along with blanks and standards, were analyzed on the ICP-MS to determine the Fe-57 concentration.

Analysis of $^{57}$Fe uptake data

To determine whether differences existed in the uptake of $^{57}$Fe between the $^{57}$Fe-amino acid treatments and the controls, we conducted a linear regression analysis. Using the SciPy 1.9.3 library, we applied the linear regression function to the P-normalized concentrations of $^{57}$Fe ([$^{57}$Fe]/[P]) to estimate the slopes, intercepts, p-values, and standard errors of the slopes. This analysis tested the null hypothesis that the slope of the regression line is zero, employing a Wald test with a t-distribution for the test statistic.

To further evaluate iron uptake enhancement mediated by the $^{57}$Fe-amino acid complexes, we analyzed the time-course data for both $^{57}$Fe and $^{56}$Fe isotopes across the different treatments. Specifically, we aimed to test whether the addition of Fe complexed to an amino acid is bioavailable by comparing the rate of iron uptake for $^{57}$Fe relative to $^{56}$Fe. If bioavailable, the rate of $^{57}$Fe should be higher than that of $^{56}$Fe. To test this, we fitted an ordinary least squares (OLS) linear model to the concentration data for each isotope over time, incorporating an effect for the isotope and its interaction with time. The model was specified as follows:

$$[\text{Fe isotope}] = \beta_0 + \beta_1 \cdot \text{Time} + \beta_2 D_{Fe57} + \beta_3 \cdot (\text{Time} \times D_{Fe57}),$$

Where the beta coefficients represent the model parameters and $D_{Fe57}$ is a dummy variable equal to 1 when the concentration modeled is that of $^{57}$Fe and 0 when it is that of $^{56}$Fe. The OLS linear model was implemented using the statsmodels 0.15.0 library and Python version 3.11.8.

**Supplementary Text 1**

Assessment of Nutrient Depletion Across Three Cell Layers in a Flow System

We evaluated whether the nutrient consumption by three monolayers of cells inside the MCCD leads to significant nutrient depletion across the layers. To quantify this effect, we introduce a dimensionless quantity $\frac{3k}{vC_o}$, which compares the rate of nutrient consumption by the cell layers to the rate at which nutrients are supplied by the flowing medium.

Derivation of the Dimensionless Quantity:
The system consists of three monolayers of cells on a membrane, over which a medium flows. The key parameters involved in the analysis are:

- Flow velocity $v$: The speed at which the medium flows across the cell layers.
- Initial nutrient concentration $C_o$: The concentration of nutrients in the medium before any consumption by the cells.
- Nutrient consumption rate $k$: The rate at which nutrients are consumed by each layer of cells per unit area.
- Number of cell layers: Three.

The flow of nutrients across the first cell layer is proportional to the product of the flow velocity and the initial nutrient concentration, $vC_o$. Meanwhile, the consumption of nutrients by each cell layer is proportional to the product of the consumption rate and the surface area, $kA$. To assess the significance of nutrient consumption across all three layers, we sum the total consumption and compare it to the nutrient supply rate:

$$\text{Total nutrient consumption} = 3kA$$

$$\text{Nutrient supply rate} = vC_oA$$

By taking the ratio of these quantities, we obtain the dimensionless quantity:

$$\frac{3kA}{vC_oA} = \frac{3k}{vC_o}$$

This dimensionless quantity allows us to assess whether nutrient consumption by the cell layers is negligible. If $\frac{3k}{vC_o} < 1$, the nutrient concentration does not drop as the medium traverses the cell layers.

Parameters:

1. Flow velocity $v$: The local flow velocity was obtained from the results of the Comsol simulation (see Fig. S3). The flow velocity was uniform across the majority of the surface and had a value of ~$316\times10^{-6}$ m/s.
2. Surface are of the filter $A$: 10 $cm^2$
3. Initial nutrient concentration $C_o$: We used the known concentrations of each amino acid and converted them to concentrations per unit area. For example, serine had a concentration of 50 µM in the High condition. With a surface area in the filter of 10 $cm^2$, this value is equivalent to 0.05 $M/m^3$.
4. Consumption rates: first, we obtained the amino acid composition of an *E. coli* cell as fractions of 1 (*81*). Then, using the cell numbers per cell layer, we calculated the amount of each amino acid per cell layer. We transformed this amount to molarity (M) and

multiplied it by the growth rate in each nutrient condition: $m_{serine} \times$ growth rate$_{\text{High condition}}$, for example. This calculation corresponds to the moles of serine consumed to produce a new cell. That is, this consumption rate assumes that amino acids are consumed to synthesize protein only and not for carbon metabolism. To contemplate both uses of degradable amino acids, uptake rates would have to be measured directly.

Results:
The values of the dimensionless quantity are presented below for each nutrient condition.

| | High | Medium | Low |
|---|---|---|---|
| L-Alanine | 0.09673991 | 0.11340039 | 0.49906263 |
| L-Arginine HCl | 0.00438288 | 0.0051377 | 0.02261044 |
| L-Asparagine | 0.06122259 | 0.07176631 | 0.31583561 |
| L-Aspartic Acid | 0.06077182 | 0.0712379 | 0.31351015 |
| L-Glutamic Acid | 0.04001219 | 0.04690306 | 0.20641521 |
| L-Glutamine | 0.04028325 | 0.0472208 | 0.20781353 |
| L-Glycine | 0.13692142 | 0.16050194 | 0.70635131 |
| L-Histidine HCl $H_2O$ | 0.04097939 | 0.04803684 | 0.21140483 |
| L-Isoleucine | 0.07432234 | 0.08712209 | 0.38341469 |
| L-Proline | 0.06442814 | 0.07552391 | 0.33237241 |
| L-Serine | 0.00275611 | 0.00323077 | 0.01421827 |
| L-Threonine | 0.07146233 | 0.08376953 | 0.36866044 |
| L-Tryptophan | 0.0373576 | 0.04379131 | 0.19272071 |
| L-Valine | 0.08080483 | 0.09472098 | 0.4168566 |
| L-Leucine | 0.05762674 | 0.06755118 | 0.2972853 |
| L-Lysine HCl | 0.07876708 | 0.0923323 | 0.40634425 |
| L-Methionine | 0.06912414 | 0.08102865 | 0.35659816 |
| L-Phenylalanine | 0.03763342 | 0.04411463 | 0.19414361 |
| L-Cysteine HCl | 0.10145298 | 0.11892514 | 0.52337643 |
| L-Tyrosine | 0.05107541 | 0.05987159 | 0.26348825 |

All the values are below 1 in all nutrient conditions. However, some of these values are close to 1, particularly those for alanine, glycine, and cysteine in the Low condition. Nonetheless, these calculations show that cells do not deplete the amino acids as the media traverses the cell layers in any of the three conditions.

**Supplementary Text 2**

<u>Oxidation of ferrous iron to ferric iron during our experimental timescales</u>

Ferrous iron (Fe(II)) is rapidly oxidized to ferric iron (Fe(III)) in oxic environments unless chelated. The added iron to our media was ferrous iron chelated with tricine, but before assuming that the total added iron remained in the ferrous state, we estimated the amount of ferrous iron that could be potentially oxidized to ferric iron.

We conservatively calculated the potential oxidation of Fe(II) during the experimental timescales (5-15 h). The oxidation rate of the sum of inorganic species, Fe(II)', using a second-order rate law and ambient oxygen concentration:

$$d\mathrm{Fe(II)}/dt = k[\mathrm{Fe(II)}][O_2].$$

This calculation yielded a value of $1\times10^{-7}$ M/day, indicating that it would take approximately 100 days to oxidize Fe(II)' to Fe(III). For this calculation, we used a Fe(II)' oxidation rate by $O_2$ of $k = 0.864\ \mu\mathrm{M}^{-1}\mathrm{d}^{-1}$ and an $O_2$ concentration of $[O_2] = 214\ \mu\mathrm{M}$ (*82*). This calculation does not account for oxygen consumption by *E. coli* or other processes (such as secretion of reducing compounds), which would further slow Fe(II) oxidation. We conclude that Fe(III) levels in our media were negligible, justifying our focus on Fe(II). Furthermore, complexation with the media constituent tricine would further lessen this oxidation

In standard culture conditions with scarce iron, siderophores produced by *E. coli* would bind Fe(III) keeping it soluble and available. However, in the MCCD these siderophore-Fe(III) complexes are continually washed out allowing us to examine the other iron uptake mechanisms.

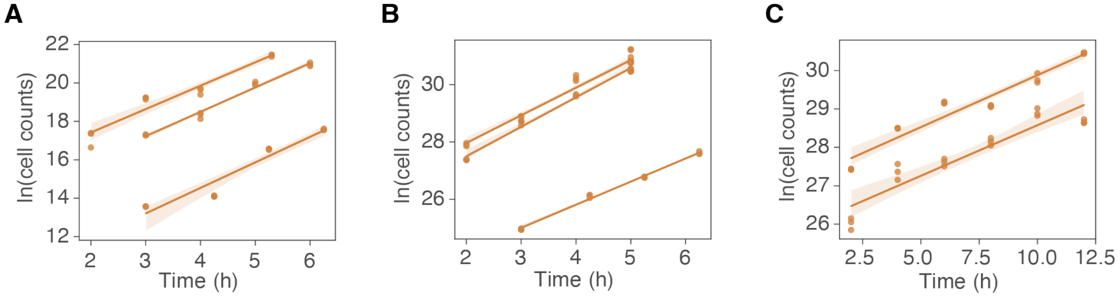


**Figure S1. Growth curves from experiments in the MCCD.** Growth experiments were performed under High **(A)**, Average **(B)**, and Low **(C)** media conditions. These growth curves were used to calculate the specific growth rates reported in Fig. 1B.

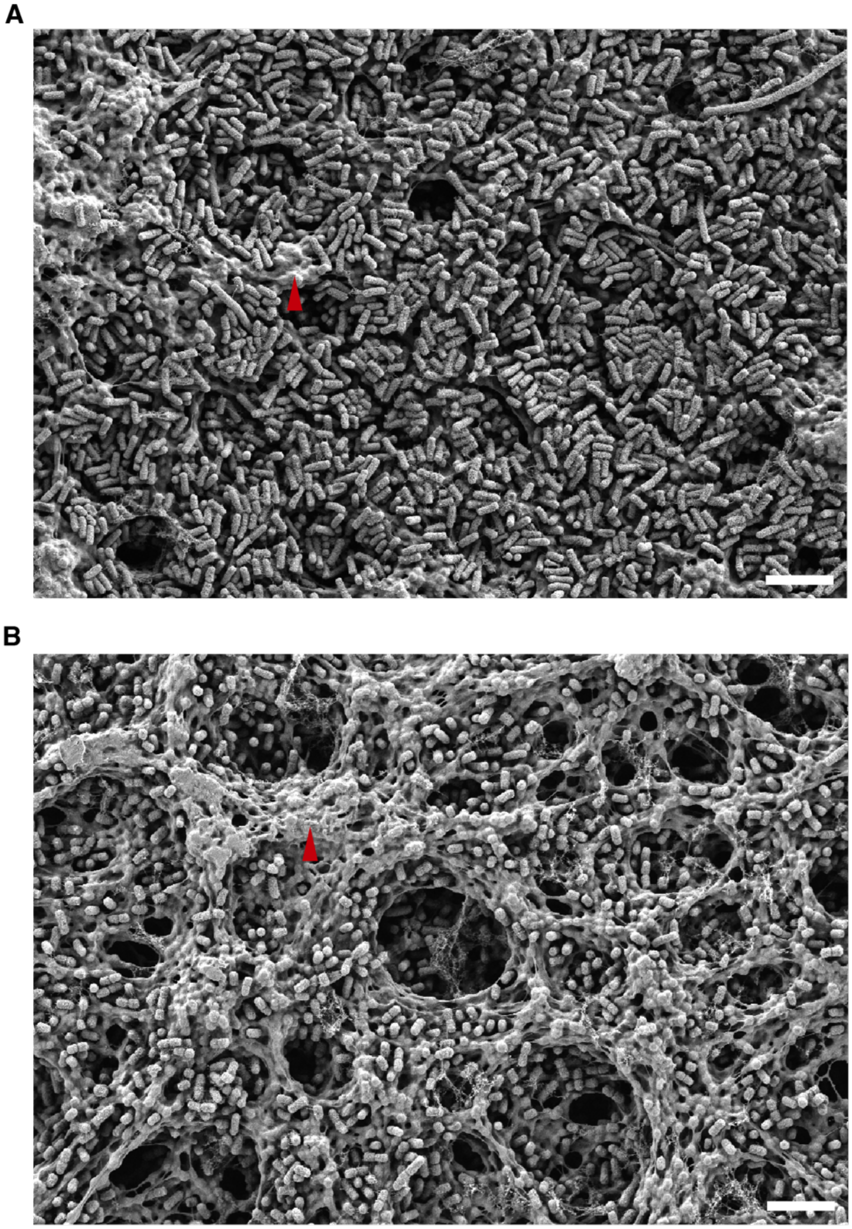


**Figure S2. Representative SEM images confirmed that there were three cell layers at most during sampling times.** Images were captured at approximately the same cell densities as the samples for proteomics. These images show at most three cell layers with no evidence of cell cluster formation in the High medium condition (**A**) and the Low medium condition (**B**). Sections of the filter membrane (red arrowheads) are clearly visible, indicating the absence of multiple stacked cell layers. Scale bars: 5 µm. Approximately 20% of the total cells in the culture at the time of sample preparation for SEM were dislodged from the filter, but these dislodged cells would account for only about 10% of a full additional cell layer on the membrane surface.

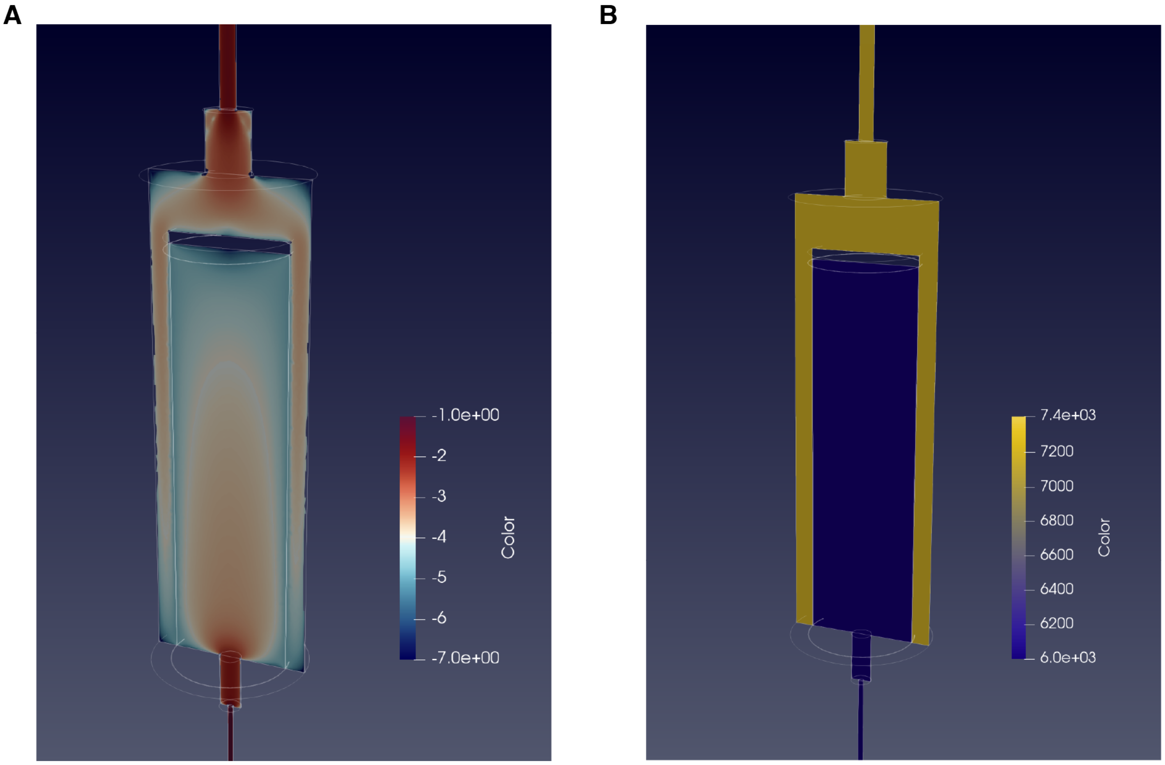


**Figure S3. Uniform flow velocity and low pressure across the filter membrane.** Numerical simulations of fluid flow through the Sterivex filter yielded values for flow velocity on the filter membrane **(A)** and the pressure exerted on the surface by the medium flow **(B)**. The color gradient in (A) represents the logarithm of the fluid velocity (m/s), and the color gradient in (B) represents the pressure (Pa) under experimental conditions. These flow velocity and pressure values were generated using the permeability values without bacteria. Changes to flow velocity and pressure with bacteria were negligible. Though the pressure is mild, visual inspections before and after unplugging the filters indicate that this pressure is sufficient for keeping the cells on the surface of the membrane. That is, cells do not grow in the space between the membrane and the plastic casing.

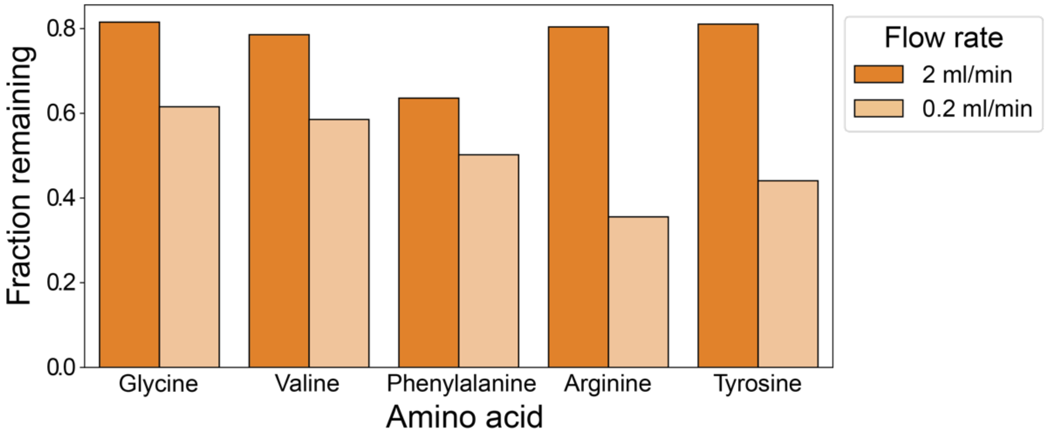


**Figure S4. Metabolomics measurements of amino acids consumed by *E. coli* in High medium.** We quantified the fraction remaining of five amino acids (glycine, valine, phenylalanine, arginine, and tyrosine) by comparing their concentrations in the outflow to their concentrations in the inflow at two flow rates: 2 ml/min and 0.2 ml/min. The results show that these amino acids were not depleted. No other amino acids were detected in the inflow or outflow, as their concentrations in the High medium were below the limit of detection. The cell counts at the time of media collection were 3.35 x $10^9$ and 4.18 x $10^9$ for the 2 ml/min and the 0.2 ml/min flow rates, respectively. These cell counts are in the same order of magnitude as the samples collected for proteomic analysis.

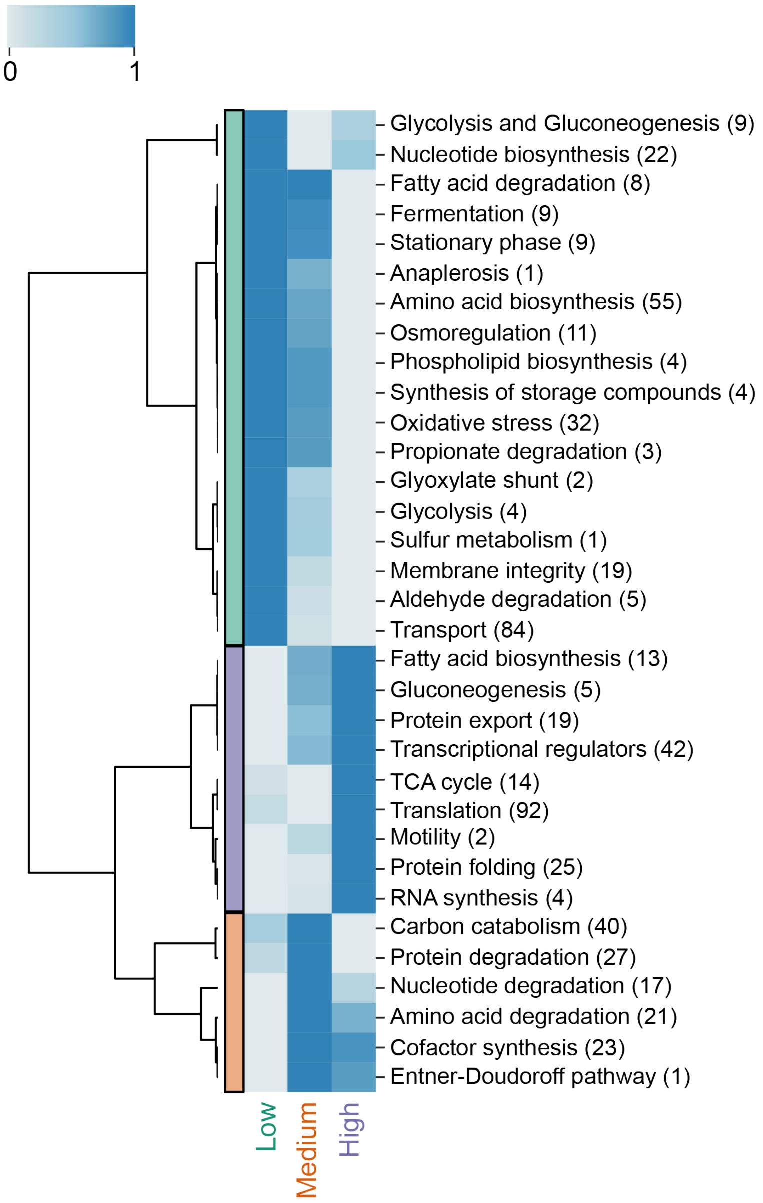


**Figure S5. Protein expression profiles of functional groups under the three nutrient conditions.** Complete heatmap displaying the expression profiles across nutrient conditions. As in Fig. 2, the heatmap groups proteins by biological function (see Materials and Methods) and sums protein abundance within each group for each replicate of the nutrient conditions. The heatmap displays the median value of these summed protein abundances. The color of the heatmap displays the median values after normalizing them across each row by assigning the lowest value 0 (lightest blue) and the highest value 1 (darkest blue). Only protein groups with statistically significant differences across conditions ($p < 0.05$, Kruskal-Wallis test). Numbers in parentheses indicate the number of proteins in each group. This figure includes protein groups with a single protein (Anaplerosis, Sulfur metabolism, and Entner-Doudoroff pathway) as well as the Transcriptional regulators group. The latter group, which contains many proteins controlling various processes, was excluded from the main figure as transcriptional regulation was not the focus of this study and warrants further investigation in future studies.

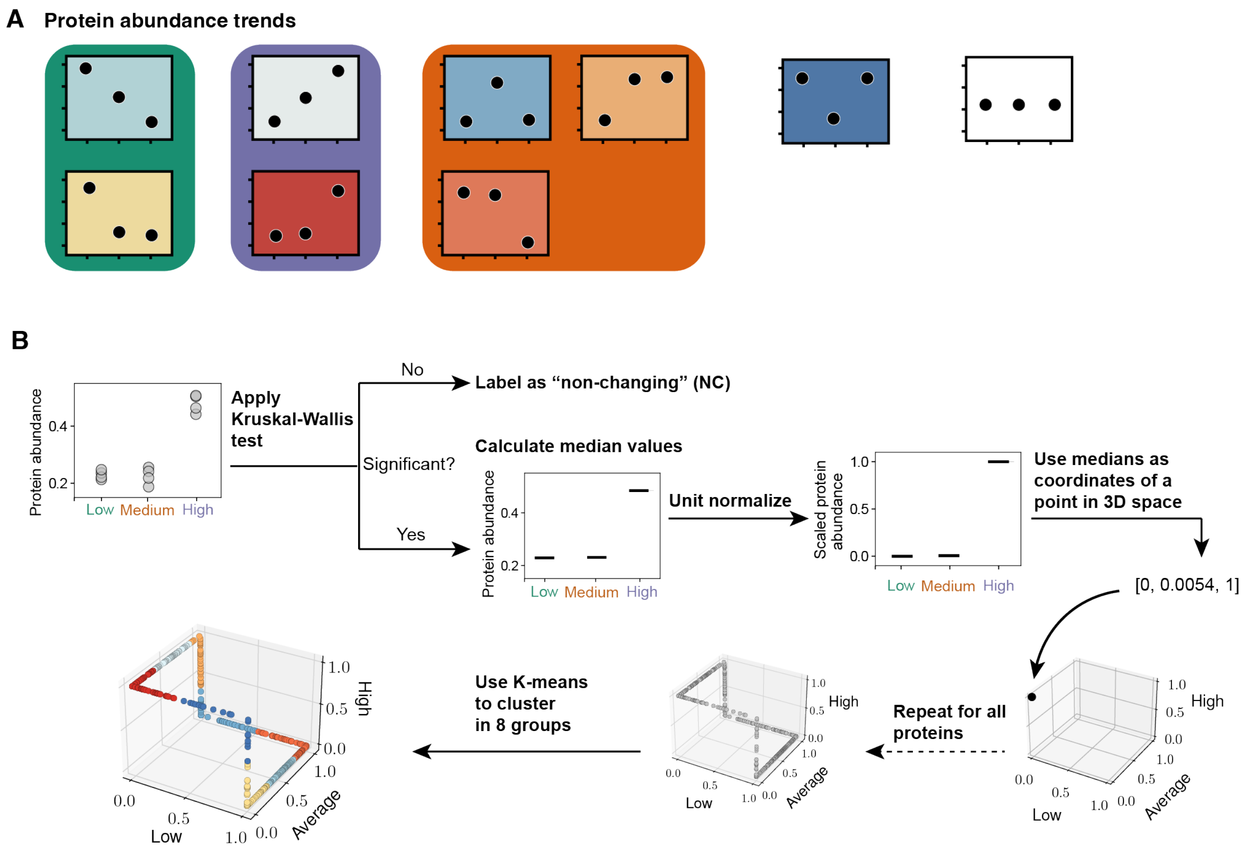


**Figure S6. Method for classifying proteins in the Carbon catabolism, Nucleotide degradation, and Proteases functional groups.**
**(A)** Visual inspection of individual protein abundance scatterplots revealed eight distinct trends across conditions. The first group (green background, corresponding to the green group in Fig. 2 and Fig. S6) shows peak protein abundance in the Low condition. The second group (purple background, corresponding to the purple group in Fig. 2 and Fig. S6) peaks in the High condition. The third group (orange background, corresponding to the orange group in Fig. 2 and Fig. S6) peaks in the Average condition or shows comparable abundance between Average and either Low or High conditions. An additional trend shows the lowest abundance in the Average condition with comparable High and Low abundances, which we did not include in any of the three main groups (dark blue plot background). The final group includes proteins without significant change (white plot background).
**(B)** Based on this visual inspection, we implemented a clustering pipeline that maps the scaled median values of each protein (using Min-max scaling to set the highest value to one and the lowest to zero) to a point in 3D space. Proteins with significant changes ($p < 0.05$, Kruskal-Wallis test) were mapped to 3D space, and a K-means clustering algorithm was used with the number of clusters set to 8 (corresponding to the eight trends, excluding unchanging proteins, NC). The final plot shows the mapped proteins colored by trend.

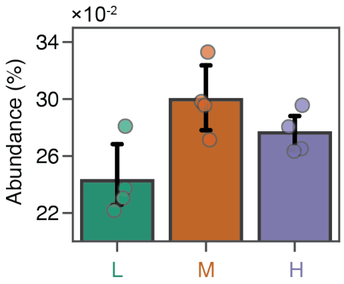


**Figure S7. Protein IscS, a subunit of the the Isc system.** In the cytoplasm, iron is used by the Isc system to assemble iron-sulfur clusters, which are essential for the catalytic activity of client proteins receiving the clusters. The barplot of the Isc system protein IscS across nutrient conditions. The barplot shows individual protein abundances for each replicate (circles), along with the mean (bar height) and the 95% confidence interval (error bars).

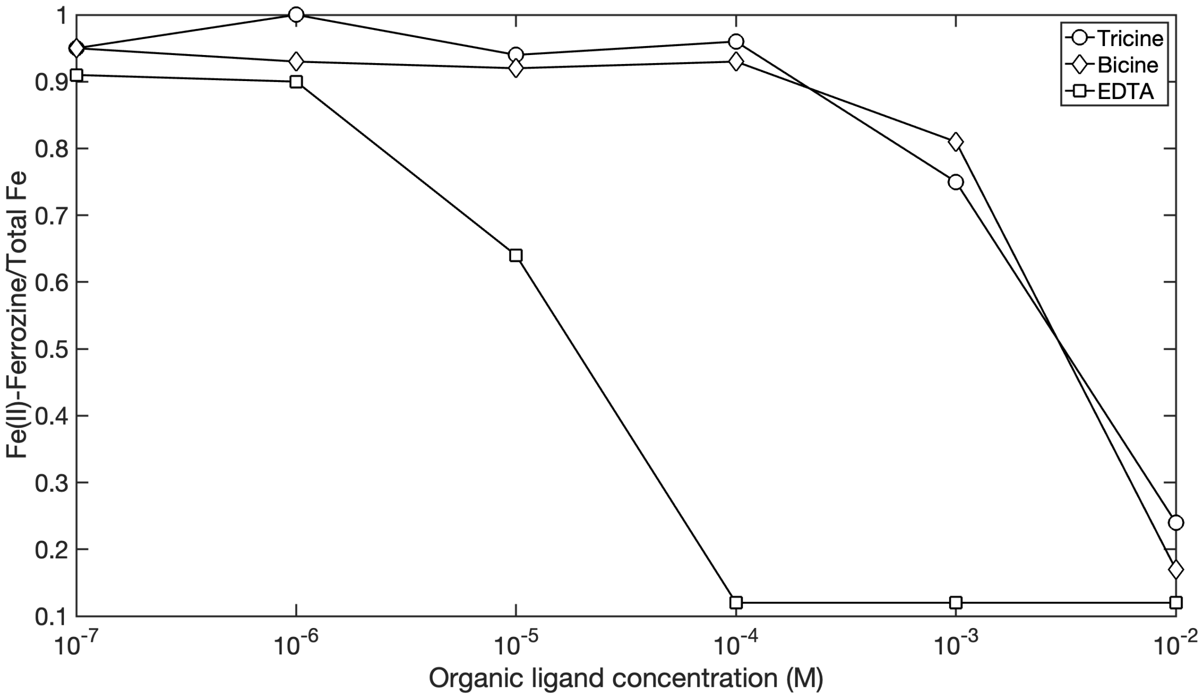


**Figure S8. Tricine and bicine exhibit similar ligand strengths.** Competitive ligand exchange between an initial addition of 10 μM Fe(II)-ferrozine and varying concentrations of tricine, bicine, and EDTA was conducted at pH 7 (sampled at equilibrium). The results demonstrate that tricine and bicine have comparable stability constants.

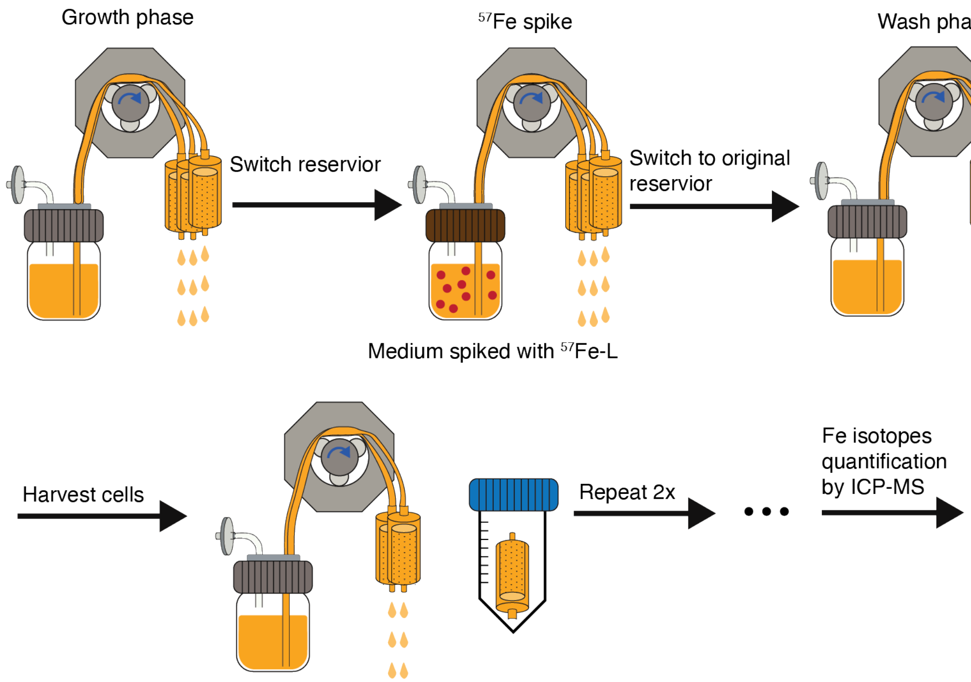


**Figure S9. Procedure for $^{57}$Fe uptake experiments.** Schematic depicting the experimental design of the $^{57}$Fe uptake experiments. First, cells were grown for 3.5 hours to reach sufficient density for downstream analyses. Second, the medium reservoir was replaced with one spiked with $^{57}$Fe complexed with a ligand. The spiked medium flowed for 10 minutes before switching back to fresh medium without $^{57}$Fe. This fresh medium flowed for another 10 minutes to wash away residual $^{57}$Fe. Cells were then harvested. The procedure was repeated twice, each time starting 30 minutes after the previous spike initiation.

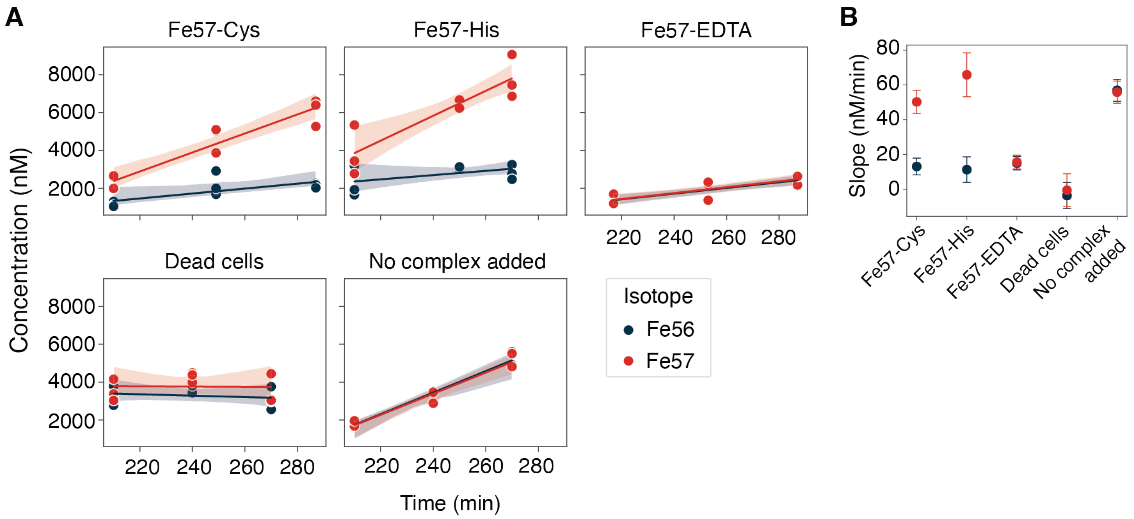


**Figure S10. Iron uptake when complexed with amino acids.** (**A**) Absolute concentrations of the iron isotopes $^{56}Fe$ and $^{57}Fe$ during the time-course experiment. Each point represents one of three biological replicates. (**B**) The average slopes of linear regression models fitted to the time-course measurements of intracellular $^{56}Fe$ and $^{57}Fe$ concentrations. Significant differences between the slopes of $^{56}Fe$ and $^{57}Fe$ for the $^{57}Fe$-Cys and $^{57}Fe$-His treatments were measured ($p < 0.05$, two-tailed t-test). The error bars represent the standard error of the estimated average slope.

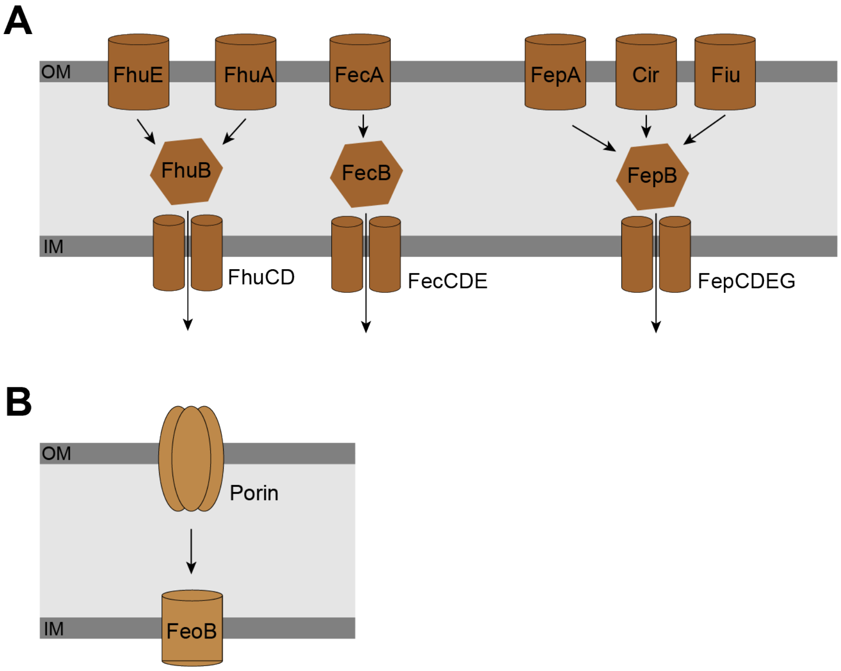


**Figure S11. Schematic of Known Iron Uptake Systems in E. coli.**
*E. coli* can acquire iron either as Fe(III) or Fe(II) through distinct pathways. **(A)** Fe(III) is primarily taken up via siderophore-mediated transport systems, where siderophores bind to Fe(III) and facilitate its transport into the cell. **(B)** Fe(II) uptake occurs via porin-mediated transport into the periplasm, followed by transport into the cytoplasm through the Feo system. Single-barrel shapes represent transporters composed of a single protein, while hexagons within the periplasm (light gray area) denote substrate-binding proteins. Shapes with two or more subunits depict multi-protein transporters.

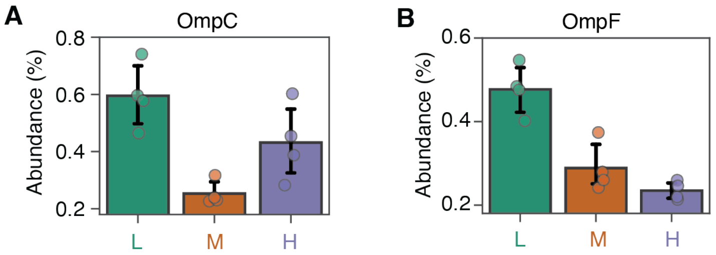


**Figure S12. Examples of transporter proteins that may be involved in iron acquisition.** Outer membrane porin OmpC (**A**) and Outer membrane porin OmpF (**B**)**.** Outer membrane porins transport small molecules such as sugars, amino acids and ions in a non-specific manner. Porins are believed to transport ferrous through the outer membrane into the periplasm.

**Supplementary Table 1.**
The three media used in this study were produced by diluting the 5x EZ and 10x ACGU solutions from the MOPS EZ Rich Defined Medium (Teknova Product number M2105) to the final concentrations listed below.

| | Concentration (µM) | | |
|---|---|---|---|
| **Nutrient** | **High medium** | **Average medium** | **Low medium** |
| Iron sulfate | 10 | 10 | 10 |
| Tricine | 400 | 400 | 400 |
| Potassium Hydroxide | 7.5000 | 3.8250 | 0.1500 |
| Adenine | 0.9950 | 0.5075 | 0.0199 |
| Cytosine | 0.9950 | 0.5075 | 0.0199 |
| Uracil | 0.9950 | 0.5075 | 0.0199 |
| Guanine | 0.9950 | 0.5075 | 0.0199 |
| L-Alanine | 4.0000 | 2.0400 | 0.0800 |
| L-Arginine HCl | 26.0000 | 13.2600 | 0.5200 |
| L-Asparagine | 2.0000 | 1.0200 | 0.0400 |
| L-Aspartic Acid, Potassium Salt | 2.0000 | 1.0200 | 0.0400 |
| L-Glutamic Acid, Potassium Salt | 3.0000 | 1.5300 | 0.0600 |
| L-Glutamine | 3.0000 | 1.5300 | 0.0600 |
| L-Glycine | 4.0000 | 2.0400 | 0.0800 |
| L-Histidine HCl $H_2O$ | 1.0000 | 0.5100 | 0.0200 |
| L-Isoleucine | 2.0000 | 1.0200 | 0.0400 |
| L-Proline | 2.0000 | 1.0200 | 0.0400 |
| L-Serine | 50.0000 | 25.5000 | 1.0000 |
| L-Threonine | 2.0000 | 1.0200 | 0.0400 |
| L-Tryptophan | 0.5000 | 0.2550 | 0.0100 |
| L-Valine | 3.0000 | 1.5300 | 0.0600 |
| L-Leucine | 4.0000 | 2.0400 | 0.0800 |
| L-Lysine HCl | 2.0000 | 1.0200 | 0.0400 |
| L-Methionine | 1.0000 | 0.5100 | 0.0200 |
| L-Phenylalanine | 2.0000 | 1.0200 | 0.0400 |
| L-Cysteine HCl | 0.5000 | 0.2550 | 0.0100 |
| L-Tyrosine | 1.0000 | 0.5100 | 0.0200 |
| Thiamine HCl | 0.0500 | 0.0255 | 0.0010 |
| Calcium Pantothenate | 0.0500 | 0.0255 | 0.0010 |
| para-Amino Benzoic Acid | 0.0500 | 0.0255 | 0.0010 |
| para-Hydroxy Benzoic Acid | 0.0500 | 0.0255 | 0.0010 |
| 2,3-diHydroxy Benzoic Acid | 0.0500 | 0.0255 | 0.0010 |

**Supplementary Table 2.**
Cell counts from each replicate sample that was used for proteomics analysis.

| Medium | Replicate | Number of cells |
|---|---|---|
| High | 1 | 1.373E+09 |
| High | 2 | 1.614E+09 |
| High | 3 | 1.608E+09 |
| High | 4 | 1.602E+09 |
| Average | 1 | 2.002E+09 |
| Average | 2 | 1.219E+09 |
| Average | 3 | 1.521E+09 |
| Average | 4 | 1.500E+09 |
| Low | 1 | 1.193E+09 |
| Low | 2 | 2.046E+09 |
| Low | 3 | 1.117E+09 |
| Low | 4 | 2.129E+09 |

**Supplementary Table S3**. **(separate file)**
Classification of the 861 proteins into functional groups. The third column ("Detail") provides more information on the biological function of each protein.

**Supplementary Table S4**. **(separate file)**

Classification of carbon catabolism, protein degradation and nucleotide catabolism proteins according to the abundance trends across conditions. See Materials and Methods and Fig. S6 for a description of the classification method. The column "Trend" indicates the condition in which the protein abundance peaks (Low, Medium or High) or if no change was detected across conditions (NC).

**Supplementary Table 5**
Stability constants used in the chemical equilibrium simulation. The list below contains stability constants for Fe2+ with amino acids and Fe2+ with bicine.

| Reaction | Log K | Enthalpy (Kcal/mol) |
|---|---|---|
| $Fe^{2+} + Ala^{-} = FeAla^{+}$ | 4.10 | |
| $Fe^{2+} + 2Ala^{-} = Fe(Ala)_2$ | 8.10 | |
| $Fe^{2+} + Arg^{-} = FeArg^{+}$ | 3.80 | |
| $Fe^{2+} + Asp^{-} = FeAsp^{+}$ | 4.80 | |
| $Fe^{2+} + 2Asp^{-} = Fe(Asp)_2$ | 9.40 | |
| $Fe^{2+} + Cys^{2-} = FeCys$ | 12.20 | |
| $Fe^{2+} + Glu^{2-} = FeGlu$ | 4.70 | |
| $Fe^{2+} + Gly^{-} = FeGly^{+}$ | 4.31 | -3.60 |
| $Fe^{2+} + 2Gly^{-} = Fe(Gly)_2$ | 8.29 | |
| $Fe^{2+} + 2His^{-} = Fe(His)_2$ | 10.90 | |
| $Fe^{2+} + His^{-} = FeHis^{+}$ | 6.30 | |
| $Fe^{2+} + Ile^{-} = FeIle^{+}$ | 3.90 | |
| $Fe^{2+} + Leu^{-} = FeLeu^{+}$ | 3.90 | |
| $Fe^{2+} + Lys^{-} = FeLys^{+}$ | 5.10 | |
| $Fe^{2+} + Met^{-} = FeMet^{+}$ | 3.70 | |
| $Fe^{2+} + 2Met^{-} = Fe(Met)_2$ | 7.50 | |
| $Fe^{2+} + Phe^{-} = FePhe^{+}$ | 3.80 | |
| $Fe^{2+} + 2Phe^{-} = Fe(Phe)_2$ | 7.10 | |
| $Fe^{2+} + Pro^{-} = FePro^{+}$ | 4.60 | |
| $Fe^{2+} + 2Pro^{-} = Fe(Pro)_2$ | 9.10 | |
| $Fe^{2+} + Ser^{-} = FeSer^{+}$ | 3.90 | |
| $Fe^{2+} + 2Ser^{-} = Fe(Ser)_2$ | 7.90 | |
| $Fe^{2+} + Thr^{-} = FeThr^{+}$ | 3.80 | |
| $Fe^{2+} + 2Tyr^{2-} = Fe(Tyr)_2^{2-}$ | 8.20 | |
| $Fe^{2+} + 2Val^{-} = Fe(Val)_2$ | 7.60 | |
| $Fe^{2+} + Val^{-} = FeVal^{+}$ | 3.90 | |
| $2Fe^{2+} + Bicine = Fe_2BicineO_4H_{13}^{4+}$ | 7.31 | |
| $Fe^{2+} + Bicine = FeBicineO_4H_{13}^{2+}$ | 4.31 | |

**Supplementary Table S6**

Results from the chemical equilibrium model for each nutrient condition. Listed are the concentrations of all Fe(II) complexes predicted to form with media components in each of the three nutrient conditions.

| Condition | Name | Concentration | LogC |
|---|---|---|---|
| High | Fe(2+) | 1.17E-07 | -6.930 |
| High | Fe(OH)3- | 1.01E-14 | -13.994 |
| High | Fe(OH)2 (aq) | 1.03E-13 | -12.988 |
| High | FeOH+ | 6.31E-10 | -9.200 |
| High | FeCl | 1.91E-09 | -8.719 |
| High | FeAla | 9.94E-12 | -11.003 |
| High | FeArg | 1.27E-10 | -9.895 |
| High | FeAsp | 3.94E-11 | -10.404 |
| High | FeCys | 2.97E-07 | -6.527 |
| High | FeGlu | 1.70E-11 | -10.770 |
| High | FeGly | 2.13E-11 | -10.671 |
| High | FeHis | 1.48E-09 | -8.830 |
| High | FeIle | 4.96E-12 | -11.305 |
| High | FeLeu | 9.92E-12 | -11.004 |
| High | FeLys | 1.00E-13 | -13.000 |
| High | FeMet | 4.88E-12 | -11.311 |
| High | FePhe | 9.80E-12 | -11.009 |
| High | FePro | 1.99E-12 | -11.702 |
| High | FeSer | 3.89E-10 | -9.410 |
| High | FeThr | 1.93E-11 | -10.715 |
| High | FeTyr | 4.75E-16 | -15.324 |
| High | FeVal | 7.44E-12 | -11.128 |
| High | FeBicine | 9.58E-06 | -5.019 |
| High | TOTAL Fe(2+) | 0.00001 | -5.000 |
| Average | Fe(2+) | 1.19E-07 | -6.924 |
| Average | Fe(OH)3- | 1.03E-14 | -13.987 |
| Average | Fe(OH)2 (aq) | 1.05E-13 | -12.981 |
| Average | FeOH+ | 6.41E-10 | -9.193 |
| Average | FeCl | 1.92E-09 | -8.716 |
| Average | FeAla | 5.15E-12 | -11.288 |
| Average | FeArg | 6.62E-11 | -10.179 |

| | | | |
|---|---|---|---|
| Average | FeAsp | 2.40E-11 | -10.619 |
| Average | FeCys | 1.53E-07 | -6.817 |
| Average | FeGlu | 8.80E-12 | -11.056 |
| Average | FeGly | 1.11E-11 | -10.957 |
| Average | FeHis | 7.65E-10 | -9.116 |
| Average | FeIle | 2.57E-12 | -11.590 |
| Average | FeLeu | 5.14E-12 | -11.289 |
| Average | FeLys | 5.19E-14 | -13.285 |
| Average | FeMet | 2.53E-12 | -11.597 |
| Average | FePhe | 5.08E-12 | -11.294 |
| Average | FePro | 1.03E-12 | -11.988 |
| Average | FeSer | 2.01E-10 | -9.696 |
| Average | FeThr | 9.99E-12 | -11.001 |
| Average | FeTyr | 1.25E-16 | -15.902 |
| Average | FeVal | 3.86E-12 | -11.414 |
| Average | FeBicine | 9.72E-06 | -5.012 |
| Average | TOTAL Fe(2+) | 0.00001 | -5.000 |
| Low | Fe(2+) | 1.21E-07 | -6.917 |
| Low | Fe(OH)3- | 1.05E-14 | -13.981 |
| Low | Fe(OH)2 (aq) | 1.06E-13 | -12.975 |
| Low | FeOH+ | 6.50E-10 | -9.187 |
| Low | FeCl | 1.97E-09 | -8.706 |
| Low | FeAla | 2.05E-13 | -12.689 |
| Low | FeArg | 2.63E-12 | -11.581 |
| Low | FeAsp | 8.12E-13 | -12.091 |
| Low | FeCys | 6.01E-09 | -8.221 |
| Low | FeGlu | 3.50E-13 | -12.456 |
| Low | FeGly | 4.39E-13 | -12.357 |
| Low | FeHis | 3.04E-11 | -10.517 |
| Low | FeIle | 1.02E-13 | -12.991 |
| Low | FeLeu | 2.04E-13 | -12.690 |
| Low | FeLys | 2.06E-15 | -14.686 |
| Low | FeMet | 1.01E-13 | -12.997 |
| Low | FePhe | 2.02E-13 | -12.695 |
| Low | FePro | 4.09E-14 | -13.388 |
| Low | FeSer | 7.97E-12 | -11.098 |

| | | | |
|---|---|---|---|
| Low | FeThr | 3.97E-13 | -12.401 |
| Low | FeTyr | 1.96E-19 | -18.709 |
| Low | FeVal | 1.53E-13 | -12.814 |
| Low | FeBicine | 9.86E-06 | -5.006 |
| Low | TOTAL Fe(2+) | 0.00001 | -5.000 |

**Supplementary Table S7**

Measured concentrations of Fe56 and Fe57 using ICP-MS. Sample names include the timepoint ("T" followed by the number), the type of sample (H: Fe57-Histidine; C: Fe57-Cysteine; E: Fe57-EDTA control; D: "dead cells" control; N: no complexes added), and the replicate (R1, R2, R3). The "Time" column indicates the time (in minutes) when the complexes were added after the start of growth in the MCCD.

| Sample Name | Time (min) | Complex added | Fe56 (nM) | Fe57 (nM) | Fe/P (µM/M) |
|---|---|---|---|---|---|
| **T1 H R1** | **210** | **Fe57-Histidine** | 3201.78 | 5342.79 | 4792.14 |
| **T1 H R2** | **210** | **Fe57-Histidine** | 1657.49 | 2776.86 | 4530.50 |
| **T1 H R3** | **210** | **Fe57-Histidine** | 1923.61 | 3441.60 | 4338.38 |
| **T2 H R1** | **250** | **Fe57-Histidine** | 3182.59 | 6615.62 | 5564.12 |
| **T2 H R2** | **250** | **Fe57-Histidine** | 3008.82 | 6668.78 | 5420.00 |
| **T2 H R3** | **250** | **Fe57-Histidine** | 3132.22 | 6231.61 | 5948.44 |
| **T3 H R1** | **270** | **Fe57-Histidine** | 3252.04 | 9063.84 | 6296.82 |
| **T3 H R2** | **270** | **Fe57-Histidine** | 2777.21 | 7456.18 | 6074.94 |
| **T3 H R3** | **270** | **Fe57-Histidine** | 2474.98 | 6864.79 | 6103.09 |
| **T1 E R1** | **217** | **Fe57-EDTA** | 1246.07 | 1266.11 | 2721.10 |
| **T1 E R2** | **217** | **Fe57-EDTA** | 1652.95 | 1692.13 | 2386.64 |
| **T1 E R3** | **217** | **Fe57-EDTA** | 1198.60 | 1197.03 | 2444.19 |
| **T2 E R1** | **253** | **Fe57-EDTA** | 1347.95 | 1372.04 | 2533.44 |
| **T2 E R2** | **253** | **Fe57-EDTA** | 2170.53 | 2196.18 | 2851.83 |
| **T2 E R3** | **253** | **Fe57-EDTA** | 2282.79 | 2325.37 | 2511.61 |
| **T3 E R1** | **287** | **Fe57-EDTA** | 2118.22 | 2172.84 | 2708.03 |
| **T3 E R2** | **287** | **Fe57-EDTA** | 2569.56 | 2628.80 | 2999.45 |
| **T3 E R3** | **287** | **Fe57-EDTA** | 2553.73 | 2629.70 | 2953.30 |
| **T1 C R1** | **210** | **Fe57-Cysteine** | 1305.77 | 2658.55 | 4821.79 |
| **T1 C R2** | **210** | **Fe57-Cysteine** | 1115.75 | 2034.83 | 4961.27 |
| **T1 C R3** | **210** | **Fe57-Cysteine** | 1047.41 | 1991.76 | 4890.48 |
| **T2 C R1** | **249** | **Fe57-Cysteine** | 1679.44 | 3870.17 | 6419.32 |
| **T2 C R2** | **249** | **Fe57-Cysteine** | 2917.47 | 5032.13 | 8471.72 |
| **T2 C R3** | **249** | **Fe57-Cysteine** | 1997.40 | 5098.01 | 6185.24 |
| **T3 C R1** | **287** | **Fe57-Cysteine** | 2254.70 | 6603.38 | 7806.22 |
| **T3 C R2** | **287** | **Fe57-Cysteine** | 2186.02 | 6397.04 | 7509.79 |
| **T3 C R3** | **287** | **Fe57-Cysteine** | 2027.43 | 5269.30 | 7740.96 |
| **T1 N R1** | **210** | **Nothing** | 10103.53 | 9901.50 | 14559.47 |

| | | | | | |
|---|---|---|---|---|---|
| **T1 N R2** | **210** | **Nothing** | 1695.71 | 1676.40 | 2250.97 |
| **T1 N R3** | **210** | **Nothing** | 1985.09 | 1956.03 | 2272.25 |
| **T2 N R1** | **240** | **Nothing** | 3890.52 | 3844.51 | 2495.71 |
| **T2 N R2** | **240** | **Nothing** | 2948.53 | 2881.11 | 2527.88 |
| **T2 N R3** | **240** | **Nothing** | 3523.98 | 3468.32 | 2302.66 |
| **T3 N R1** | **270** | **Nothing** | 5405.08 | 5316.03 | 2496.29 |
| **T3 N R2** | **270** | **Nothing** | 4906.85 | 4814.58 | 2458.00 |
| **T3 N R3** | **270** | **Nothing** | 5596.69 | 5508.96 | 2371.67 |
| **T1 D R1** | **210** | **Fe57-Cysteine** | 3066.68 | 3376.34 | 2477.20 |
| **T1 D R2** | **210** | **Fe57-Cysteine** | 3787.50 | 4150.56 | 2166.22 |
| **T1 D R3** | **210** | **Fe57-Cysteine** | 2773.18 | 3030.98 | 1927.28 |
| **T2 D R1** | **240** | **Fe57-Cysteine** | 3706.40 | 4489.37 | 1899.27 |
| **T2 D R2** | **240** | **Fe57-Cysteine** | 3436.15 | 3982.33 | 1889.90 |
| **T2 D R3** | **240** | **Fe57-Cysteine** | 3804.98 | 4379.19 | 2263.79 |
| **T3 D R1** | **270** | **Fe57-Cysteine** | 3756.94 | 4438.25 | 1941.38 |
| **T3 D R2** | **270** | **Fe57-Cysteine** | 2655.36 | 2978.76 | 2888.81 |
| **T3 D R3** | **270** | **Fe57-Cysteine** | 2553.45 | 3035.28 | 1976.74 |

**Supplementary Table S8**

Quantification of $^{57}Fe$ in the solutions prepared for spiking during the uptake experiments. Each solution was used across all replicates. $^{56}Fe$ concentrations are not shown as they were close to the blank values, confirming the absence of $^{56}Fe$ in these solutions.

| **Complex** | **Treatment** | **[57Fe] (mM)** |
|---|---|---|
| Fe57-Histidine | Histidine | 0.075 |
| Fe57-EDTA | EDTA | 0.079 |
| Fe57-Cysteine | Cysteine | 0.094 |
| Fe57-Cysteine | Dead cells | 0.084 |

**Data S1. (separate file)**

The mass spectrometry data has been deposited to the ProteomeXchange Consortium via the PRIDE partner repository with the dataset identifiers PXD059988 and https://doi.org/110.6019/PXD059988.
